# Deficiency of mitophagy mediator Parkin in aortic smooth muscle cells exacerbates abdominal aortic aneurysm

**DOI:** 10.1101/2024.10.30.621201

**Authors:** Hui Wang, Matthew Kazaleh, Rachel Gioscia-Ryan, Jessica Millar, Gerard Temprano-Sagrera, Sherri Wood, Fran Van Den Bergh, Muriel G. Blin, Kathleen M. Wragg, Adrian Luna, Robert B. Hawkins, Scott A. Soleimanpour, Maria Sabater-Lleal, Chang Shu, Daniel A. Beard, Gorav Ailawadi, Jane C. Deng, Daniel R. Goldstein, Morgan Salmon

**Author notes:** Corresponding author: Morgan Salmon, 2800 Plymouth Road, Ann Arbor, Michigan 48109 USA.

## Abstract

Abdominal aortic aneurysms (AAAs) are a degenerative aortic disease and associated with hallmarks of aging, such as mitophagy. Despite this, the exact associations among mitophagy, aging, and AAA progression remain unknown. In our study, gene expression analysis of human AAA tissue revealed downregulation of mitophagy pathways, mitochondrial structure, and function-related proteins. Human proteomic analyses identified decreased levels of mitophagy mediators PINK1 and Parkin. Aged mice and, separately, a murine AAA model showed reduced mitophagy in aortic vascular smooth muscle cells (VSMCs) and PINK1 and Parkin expression. Parkin knockdown in VSMCs aggravated AAA dilation in murine models, with elevated mitochondrial ROS and impaired mitochondrial function. Importantly, inhibiting USP30, an antagonist of the PINK1/Parkin pathway, increased mitophagy in VSMCs, improved mitochondrial function, and reduced AAA incidence and growth. Our study elucidates a critical mechanism that proposes AAAs as an age-associated disease with altered mitophagy, introducing new potential therapeutic approaches.

## 1. Introduction

An abdominal aortic aneurysm (AAA) is defined as a local enlargement of the infrarenal aorta, with a global prevalence of 0.92% in individuals aged 30-79, which then escalates to 1.2%-3.3% in individuals over 60 years old [1, 2]. AAAs are currently understood to be an irreversible and progressive aortic dilation with the potential to rupture, which carries a mortality rate over 81%[3]. Other risk factors for AAAs include smoking, genetics, and sex. Men are at four times the risk of developing AAA compared to women, while women experience faster aneurysm growth and have a fourfold higher risk of rupture[1]. Currently, effective clinical management relies on open surgical repair or less invasive endovascular aneurysm repair (EVAR)[4]. Clinical trials investigating the potential of statins, antihypertensives, doxycycline, and anti-inflammatory medications as alternatives or adjuncts to surgical or endovascular treatment have yet to yield definitive efficacy[5–9]. Therefore, studies exploring the pathogenesis of AAAs, identifying potential new pharmacologic targets, are crucial for advancing current clinical management.

Vascular smooth muscle cells (VSMCs) constitute the predominant cellular population within the aortic wall, playing a pivotal role in the pathogenesis of AAAs. Their contribution to AAA development is characterized by a loss of contractile phenotype, increased apoptosis, immune cell recruitment, extracellular matrix remodeling, and enhanced oxidative stress[10]. Additionally, these adverse alterations are more pronounced in aged VSMCs and could describe potential mechanisms for the increased susceptibility of elderly populations to AAAs[11, 12]. However, definitive studies investigating the causal role of aging in AAA progression currently remain unexplored and largely unknown.

Emerging evidence indicates that mitochondrial dysfunction and aberrant mitochondrial dynamics are associated with the pathogenesis of AAA. For example, damaged mitochondria release double-stranded mitochondrial DNA (mtDNA) into the cytosol, stimulating the downstream cGAS-STING, a critical innate immune pathway, promoting AAA development[13]. Separate studies demonstrated that mitochondrial fission, regulated by DRP1, is significantly increased in AAA tissues, and inhibition of DRP1 mitigates AAA progression[14]. Finally, decreased mitochondrial biogenesis, regulated by PGC1-α, has been identified in human and mouse aneurysm tissues when compared to health vascular tissue[15]. These studies suggest that the processes that regulate mitochondrial turnover and dynamics may be important in AAA progression. As a part of mitochondrial dynamics, mitophagy, a macro-autophagy that selectively degrades damaged mitochondria, is essential for mitochondrial quality control and hemostasis[16]. Recent evidence suggests that mitophagy mitigates inflammation triggered by the activation of the cGAS-STING pathway by mtDNA[17]. Given that reduced mitophagy is a hallmark of aging, and considering AAA’s prevalence in older populations, investigating mitophagy’s impact on AAA progression could represent a promising avenue for further understanding mechanisms of AAA formation or rupture[18, 19].

Previous studies, especially in Parkinson’s disease, have investigated the ubiquitin-mediated mitophagy pathway facilitated by mitochondrial kinase PINK1 and cytosolic E3 ubiquitin ligase Parkin [20, 21]. In detail, PINK1 accumulates on the outer mitochondrial membrane in damaged mitochondria with reduced membrane potential, where it recruits and phosphorylates Parkin. Activated Parkin ubiquitinates various mitochondrial membrane proteins, which are then recognized by autophagy receptors. These receptors facilitate the recruitment of LC3-decorated phagophores, leading to the encapsulation of the damaged mitochondria and subsequent fusion with lysosomes for degradation [22]. Mutations in *PINK1* or *PARK2* genes are associate with defective mitochondrial dysfunction and numerous studies suggest that targeting the PINK1/Parkin pathway could be effective in preventing and treating cardiovascular diseases [23, 24]. However, its role in the pathogenesis of AAAs has never been explored. USP30 is a deubiquitinase localized to the mitochondrial outer membrane which antagonizes PINK1/Parkin mediated mitophagy [25]. Inhibition of USP30 has been shown to enhance PINK1/Parkin-mediated mitophagy [26]. Given the availability of a USP30 inhibitor, VB-08, exploring the role of PINK1/Parkin-mediated mitophagy in the development of AAAs holds promise for identifying potential pharmacological targets.

In this study, we tested our hypothesis that Parkin-mediated mitophagy is a critical factor in the pathogenesis of AAAs. Our results demonstrate that genes related to the mitophagy pathway, as well as genes involved in mitochondrial structure and function, were downregulated in human AAA samples, coinciding with decreased protein expression of PINK1 and Parkin. Additionally, we observed reduced mitophagy in VSMCs in advanced-aged murine aortas and in young murine AAAs, both associated with decreased PINK1 and Parkin expression. Furthermore, Parkin knockout in VSMCs resulted in more severe AAA rupture and increased aortic diameter, whereas administration of a USP30 inhibitor, VB-08, reduced AAA incidence and diameter. Our findings suggest that PINK1/Parkin-mediated mitophagy represents a potential novel therapeutic target for AAA treatment.

## 2. Material and Methods

### 2.1 Ethics statement

Mice were housed and maintained at 70°F, 50% humidity, in 12-hour light-dark cycles per institutional animal protocols. Animal care and use were in accordance with the Guide for the Care and Use of Laboratory Animals. The animal protocol was approved by the University of Michigan Institutional Animal Care and Use Committee (PRO00010704) in compliance with the Office of Laboratory Animal Welfare. The authors declare that all supporting data are available within the article and its online supplementary files but can provide additional information upon request to corresponding author.

### 2.2 Mice

WT young (8-12 weeks) and aged (18-20 months) male C57BL/6J mice were obtained from The Jackson Laboratory (stock #000664). mito-QC mice, mitophagy reporter mice on the C57BL/6J background were generously provided by Dr. Ian Ganley and originally generated by Taconic Artemis[27]. These mice were bred and aged to the indicated age under specific pathogen free conditions in the animal facility at the University of Michigan. C57BL/6J Parkinfl/fl mice were a gift from the lab of Dr. Scott Soleimanpour, originally generated by Dr. Ted Dawson and Lexicon Genetics, with loxP sites flanking exon 7 of the Prkn allele [28]. C57BL/6J *Parkin^fl/fl^* mice were bred to *Myh11-cre/ER^T2^* (stock #019079, C57BL/6 background), which were purchased from The Jacson Laboratory. 6-10 weeks *Parkin^fl/fl^ Myh11-cre/ER^T2^* mice received five daily intraperitoneal injections (i.p.) of either tamoxifen, to induce gene deletion, or vehicle (corn oil) and then rested for nine days. The outcomes of *Parkin^fl/fl^ Myh11-cre/ER^T2^* mice were evaluated alongside their vehicle-treated litter-mate controls. We used male mice to for conduct the research, as a significant body of studies has focused on males in AAA. This choice is supported by two key reasons. First, like humans, male mice exhibit a four-fold higher prevalence of AAA when infused with Angiotensin II compared to females [29], offering consistent and reproducible results with an optimal sample size. Second, the widely used *Myh11-cre/ER^T2^* is carried on the Y chromosome and effective for gene deletion only in male mice [30]. All the mice were maintained on a 12-h light-dark cycle and had unrestricted access to food and water.

### 2.3 Angiotensin II infusion AAA model

We employed an abdominal aortic aneurysm model combined with a PCSK9 gain-of-function mutation with Angiotensin II infusion [31–34]. The PCSK9 gain of function mutation is induced using a single intraperitoneal injection of recombinant adeno-associated virus 8-D377Y-murine PCSK9 (PCSK9-AAV) at a dose of 5.0×10^9^ vector genomes per gram body weight purchased from the University of Pennsylvania Vector core. The mice were then fed ad libitum a high-fat diet (42% calories from fat, HFD, Teklad, catalog #88137) for 6 weeks. Starting on the third week, mice received subcutaneous infusion of Angiotensin II (Sigma, catalog #A9525) or saline at a rate of 1000 ng/kg/min using mini osmotic pumps (Alzet, model #1004) for the subsequent four weeks [35].

### 2.4 Elastase application AAA model

As a secondary model for AAA, periadventitial elastase was applied following established methods[36, 37]. Male *Parkinfl/fl Myh11-cre/ER^T2^* mice received a series of injections of either corn oil (control, n=10/group) or tamoxifen (n=11/group) at 6-8 weeks of age. One month following injection, the mice underwent a midline laparotomy to expose the infra-renal aorta, which was then treated with full-strength elastase (Sigma, catalog #E1250). Each surgeon determined the optimal elastase dose and duration through a pre-experimental dose-response curve, reliably producing a 110% increase in aortic diameter in control mice. Fourteen days post-operatively, the infra-renal abdominal aortas were harvested from both groups. During the harvest, the abdominal aorta was exposed similarly, and maximal aortic diameters were measured using video micrometry. To calculate the percentage of diameter dilation, a specific section of the aorta not exposed to elastase served as an internal control.

### 2.5 USP30 inhibitor studies

For C57/B6 male mice (n=25/group) receiving either VB-08 (25 mg/kg, i.p., daily, Vincerebio.com) or a vehicle (confidential, i.p., daily, Vincerebio.com). Treatment began nine days after the initiation of PCSK9 AAV and a HFD, and five days before the implantation of an Angiotensin II pump. The treatment continued for the following 5 days and 4 weeks during the induction of abdominal aortic aneurysm (AAA), or until the mice succumbed to aneurysm rupture within this period. Mice were harvested, and anatomical pictures were taken. Tissues were processed according to experimental purposes.

### 2.6 Cholesterol and blood pressure measurements

Serum cholesterol levels in fasting mice were determined using a colorimetric assay (Cell Biolabs, catalog #STA-384). Blood pressure in mice was obtained using the CODA® non-invasive blood pressure system (Kent Scientific Corporation) via the tail-cuff method.

### 2.7 Aortic aneurysm assessments (rupture and diameters)

Following the implantation of the Angiotensin II pump, autopsies were conducted on mice that were found deceased. Deaths due to aortic aneurysm rupture were noted for inclusion in survival curve analysis. Four weeks post-implantation, mice were euthanized with CO₂ and a thoracotomy was performed. Blood was collected from the right ventricle, and the mice were perfused with cold PBS through the left ventricle after opening the right atrium. The abdominal cavity was then opened, and visceral organs were removed. The fat tissue and connections around the entire aorta and adjacent heart were isolated under a dissecting microscope. Aortas with aneurysms were identified under the microscope, and the corresponding mice were documented. The entire murine aorta and adjacent heart were isolated and pinned to the wax-coated base of a Petri dish alongside a ruler for scale. Photographs were taken, and the absolute value of maximal diameters of the aortic aneurysms were measured using ImageJ software (National Institutes of Health, USA).

### 2.8 Isolation and culture of primary aortic VSMCS

Vascular smooth muscle cells (VSMCs) were extracted from the abdominal aortas of 5-week-old male mito-QC mice using an enzymatic digestion method. Following euthanasia, the aorta was dissected under sterile conditions. The fatty and adventitial layers were removed through an 8–10-minute digestion with an enzyme cocktail consisting of 1 mg/ml collagenase type II (Worthington Biochemical Corp, catalog #LS004176), 1 mg/ml soybean trypsin inhibitor (Worthington Biochemical Corp, catalog #LS003570), and 0.744 units/ml elastase (Worthington Biochemical Corp, catalog #LS002279), all at 37°C in a 5% CO2 atmosphere. The aortas were then longitudinally sliced open and rinsed three times to remove the endothelial cell layer. These aortas were then segmented into 1-2 mm pieces and incubated in a fresh enzyme cocktail at 37°C in 5% CO2 for one hour. Post centrifugation, the resulting pellet was resuspended and cultured in VSMC media composed of 10% FBS (Gibco, catalog #16000044), 100 unit/ml penicillin and 100ug/ml streptomycin (Gibco, catalog #15140122) in DMEM/F12(Gibco, catalog #11320033). The media was refreshed every 2-3 days, and cells were passaged upon reaching approximately 80% confluence. Cells were utilized up to the eighth passage.

### 2.9 Cell culture, pharmacological intervention and Lentiviral shRNA transduction

MOVAS (ATCC, catalog #CRL-2797) were cultured in DMEM/F12 (Gibco, catalog #11320033) supplemented with 10% FBS (Gibco, catalog #16000044), 100 unit/ml penicillin and 100ug/ml streptomycin (Gibco, catalog #15140122) at 37°C in 5% CO2. Cells were passaged at 80% confluence. MOVAS and VSMCs were starved in DMEM/F12 containing 0.2% FBS for 48 hours and then received 100nM, 500nM or 1000nM Angiotensin II (Sigma, catalog #A9525) for an additional 48 hours. The USP30 inhibitor was administered at concentrations of 100nM, 200nM, 500nM or 1000nM, either concurrently with or 24 hours prior to the Angiotensin II treatment. Particularly, to assess mitophagy level changes in MOVAS cells, Mtphagy dye (Dojindo, catalog # MT02-10) was incubated for 30 minutes prior to any pharmacological intervention, in accordance with the manufacturer’s protocol.

Lentiviral shRNA targeting Parkin (Santa Cruz, catalog #sc-42159-V) and control shRNA particles (Santa Cruz, catalog #sc-108080) were purchased from Santa Cruz Biotechnology. Transduction and selection of transduced cells were performed according to the manufacturer’s protocol. Briefly, MOVAS cells were seeded in 12-well plates and cultured to 70-80% confluence. Cells were then transduced with Parkin shRNA or control lentiviral particles at a multiplicity of infection (MOI) of 0.25, 0.5, or 1, in the presence of 5 µg/mL polybrene (Santa Cruz, catalog #sc-134220). Following overnight incubation, the medium was replaced with fresh complete media, and cells were incubated for an additional 24 hours. To select stably transduced cells, the cultures were split and maintained in complete medium supplemented with 2 µg/mL puromycin (Santa Cruz, catalog #sc-108071) for 7 days, with medium changes every 2-3 days. Surviving cells were then expanded for further experimentation. Knockdown efficiency was confirmed by Western blot analysis.

### 2.10 MitoSOX Red, MitoTracker, and TMRM staining in cells

After Angiotensin II treatment, cells were detached using 0.05% Trypsin-EDTA (Gibco, 25300054) and collected. LIVE/DEAD^TM^ Aqua (Thermo Scientific, catalog #L34957) staining was performed for 5 minutes. Subsequently, cells were incubated with 5µM MitoSOX Red (Thermo Scientific, catalog #M36008) or 25nM MitoTracker Deep Red (Thermo Scientific, catalog #M22426) along with 100nM TMRM (Thermo Scientific, catalog #I34361) in a dark environment at 37°C. Following incubation, cells were washed, and flow cytometry acquisition using a ZE5 Cell Analyzer was conducted immediately.

### 2.11 Flow cytometry

For the cellular samples requiring staining with MitoSOX Red, MitoTracker or TMRM, cells were processed and stained according to method *2.10*. Subsequently, flow cytometric analysis was then conducted using a ZE5 Cell Analyzer (Bio-Rad). For the murine aortic samples processing, aortas were isolated following PBS perfusion and then slightly sectioned longitudinally into strips. These tissue strips were then enzymatically digested into single-cell suspensions using a cocktail containing 3.2mg/ml collagenase Type II (Worthington Biochemical Corp, catalog #LS004176), 0.7mg/ml elastase (Worthington Biochemical Corp, catalog #LS002279), 0.2mg/ml soybean trypsin inhibitor (Worthington Biochemical Corp, catalog #LS003570) in RPMI 1640(Gibco, catalog #11875093) at 37°C. The digestion took 30 minutes and was facilitated by intermittent pipetting. The resulting cell suspension was filtered through a 70µm mesh. Red blood cells were lysed using a 2-minute incubation with RBC lysis buffer (BioLegend, catalog #420302) and the cells were then washed. For staining, cells were first labeled with LIVE/DEAD^TM^ Aqua for viability assessment for 5 minutes, followed by a 5-minute incubation with Fc blocking reagent, anti-mouse CD16/32 (BioLegend, catalog #101320). A cocktail of surface antibodies was then applied for 30 minutes, including Brilliant Violet 785™ CD31 antibody (BioLegend, catalog #102435), APC CD90.2 (Thy-1.2) antibody (BioLegend, catalog #140312) and Alexa Fluor® 700 CD45 antibody (BioLegend, catalog #103128). After surface staining, cells were fixed, permeabilized using BD Cytofix/Cytoperm™ (BD Biosciences, catalog #555028), and stained with intracellular antibody for α-actin (Invitrogen, catalog #14-9760-82) and subsequent secondary antibody (BioLegend, catalog #405317) for 30 minutes each. Flow cytometric data acquisition was again performed on a ZE5 Cell Analyzer (Bio-Rad). All the data analysis was conducted by FowJo^TM^ Software_v10.9.0 (Ashland, 2023).

### 2.12 Western blotting of MOVAS and aortic tissues

Cellular and aortic tissue proteins were extracted using RIPA buffer (Sigma-Aldrich, catalog #R0278) supplemented with 1% protease inhibitor (Sigma-Aldrich, catalog #P8340) and phosphatase inhibitor (Sigma-Aldrich, catalog #P5726). Protein concentration was determined by the BCA protein assay per manufacturer’s protocol (Thermo Scientific, catalog #23225). Equal amounts of protein (20-25mg) were separated onto a 10% Nupage SDS-PAGE gels (Thermo Scientific, catalog #NP0315BOX, NP0303BOX) and then transferred to PVDF membranes (Thermo Scientific, catalog #IB24001) through semi-dry method. Membranes were blocked in 5% BSA (Gemini Bio-Products, catalog #700-100P) in 0.1% TBST for 1 hour at room temperature, then incubated with primary antibodies overnight at 4°C on a shaker. After washing, the membranes were incubated with HRP-conjugated secondary antibodies (abcam, catalog #ab205718, ab205719) for 1 hour at room temperature. Bands were visualized using enhanced chemiluminescence (Thermo Scientific, catalog #34580, 34094) and captured with ChemiDoc MP imaging system (Bio-Rad). Band intensity was quantified with ImageJ software and normalized to Vinculin, β-tubulin or GAPDH. All the antibodies used here are listed in supplementary table 1.

### 2.13 Frozen section and mitophagy measurement of GFP and mCherry fluorescence

Aortic segments were harvested and immersed in 3.7% paraformaldehyde within 200 mM HEPES buffer for overnight fixation. Subsequently, the fixed tissues were cryo-preserved in OCT compound (Sakura, catalog #4583) and sectioned into 6 µm slices using a Leica Cryostat (model CM1850-3-1). The sections were then placed on slides and sealed with Prolong Diamond mounting media (Thermo Scientific, catalog #P36961). Imaging was conducted with a Nikon A1si confocal microscope, utilizing a 100x magnification objective lens across both GFP and mCherry channels. Quantitative analysis of mCherry puncta was performed using ImageJ software.

### 2.14 qPCR analysis

RNA extraction from aortic tissue was conducted using the TRIzol-ethanol method. The concentration of extracted RNA was determined using a NanoDrop spectrophotometer. Subsequently, 1 µg of RNA was reverse transcribed into cDNA using the iScript cDNA Synthesis Kit (Bio-Rad, catalog #1708891). TaqMan primers, obtained from Thermo Fisher Scientific, were employed for qPCR, which was performed according to the TaqMan Advanced Master Mix Kit protocol (Applied Biosystems, catalog #4444557) on a QuantStudio 3.0 Real-Time PCR System.

Gene expression levels were quantified based on the fold change in Ct values. The details of the primers used are listed in supplementary table 1.

### 2.15 Immunohistochemistry and immunofluorescence

Aortic segments were harvested and immersed in a 4% paraformaldehyde buffer for overnight fixation. Subsequently, the fixed tissues were dehydrated, cleared, embedded, sectioned, and mounted on positive-charged slides. For the staining, tissue sections were deparaffinized and hydrated. Endogenous peroxidase activity was quenched by hydrogen peroxide in methanol for 30 minutes. Antigen retrieval was performed using citrate buffer (pH 6.0, Vector, catalog #H-3300-250) in a microwave for 20 minutes, followed by blocking non-specific binding with 10% horse serum at room temperature. Supplementary table 2 lists all the antibodies used in the subsequent steps. For immunohistochemistry, primary antibody was applied to the sections and incubated overnight at 4°C. After washing by PBS, secondary antibody was applied to the sections and incubated for 1 hour at room temperature. The signal was detected using DAB for 5-10 minutes. Then the sections were counterstained by hematoxylin, dehydrated, cleared, and mounted with Permount^TM^ mounting media (Fisher Scientific, catalog #SP15-100). Images were captured and quantification of staining intensity was performed using ImageJ software. For immunofluorescence, primary antibodies conjugated with distinct fluorophores were applied to the sections and incubated overnight at 4°C. After washing by PBS, sections were stained by DAPI. These sections were then placed on slides and sealed with Prolong Diamond mounting media (Thermo Scientific, catalog #P36961). Imaging was conducted with a Nikon A1si confocal microscope, utilizing a 60x magnification objective lens across channels of DAPI, Alexa Fluor 488, 555 and 647.

### 2.16 MitoSox Red staining for tissue

Aortic segments were excised and immediately snap-frozen in OCT compound (Sakura, catalog #4583). The blocks were sectioned to a thickness of 6 µm and mounted onto slides. Sections were rinsed with HBSS, followed by incubation in 5 µM MitoSox Red (Thermo Scientific, catalog #M36008) for 1 hour at 37°C. After a subsequent wash with HBSS, slides were promptly imaged using a Nikon A1si confocal microscope with a 100x magnification objective lens.

### 2.17 Oxygraph analysis of murine aortas

Aortas were isolated and maintained in BIOPS buffer on ice, followed by permeabilization with 50 µg/ml saponin for 25 minutes, and subsequently rinsed in MiR05 for 10 minutes [38]. The wet weights of the aortas were then blotted, measured, and documented. High-resolution measurements of oxygen consumption in permeabilized aortic tissue were performed in 2 ml of MiR05 buffer using the Oroboros Oxygraph-2k (O2k; Oroboros Instruments, Innsbruck, AT). The oxygen consumption rate was determined from plateau values on the oxygraphy chart and expressed as pmol s^-1^ per mg of wet weight. All measurements of respiration were carried out at 37 °C within an oxygen concentration working range of approximately 190-50 µM. As published previously by our lab [39], the respiration process was assessed through sequential titrations of substrates and inhibitors: 11.25 mM ADP (CalBiochem, catalog #117105), 0.5 mM malate (Sigma, catalog #M1000), 5 mM pyruvate (Sigma, catalog #P2256), 10 mM glutamate (Sigma, catalog #G1626), 10 mM succinate (Sigma, catalog #S2378, mentioned twice and may need correction), 0.5 µM CCCP (Sigma, catalog #C2759), 0.5 µM rotenone (Sigma, catalog #R8875), and 2.5 µM antimycin-A (Sigma, catalog #A8674).

### 2.18 Mitochondrial DNA copy numbers

Relative mtDNA copy numbers by qPCR with SYBR-based detection (Thermo scientific, catalog #A46109) was conducted following DNA isolation with the Blood/Tissue DNeasy kit (Qiagen, catalog #69504) according to manufacturer’s protocols. Primer sequences: mt9/mt11 (5′-GAGCATCTTATCCACGCTTCC-3′ and 5′-GGTGGTACTCCCGCTGTAAA-3′) for mtDNA and Ndufv1 (5′-CTTCCCCACTGGCCTCAAG-3′ and 5′-CCAAAACCCAGTGATCCAGC-3′) for nuclear DNA. The fold changes in mt9/mt11 expression were normalized to the Ndufv1 gene [40].

### 2.19 Metabolomics experiments and analysis

Targeted metabolomics analyses were performed as reported by Seals Laboratory and carried out by the University of Colorado Metabolomics Core Facility[41], plasma (20 µL) were extracted in 1 mL of ice-cold lysis/extraction buffer (methanol:acetonitrile:water 5:3:2). After discarding protein pellets, water and methanol soluble fractions were injected into a C18 reversed phase column (phase A: water, 0.1% formic acid; B: acetonitrile, 0.1% formic acid - Phenomenex) through an ultra-high performance chromatographic system (UHPLC, Ultimate 3000; Thermo Fisher). UHPLC was coupled online with a high-resolution quadrupole Orbitrap instrument run in either polarity modes (QExactive, Thermo Fisher) at 70,000 resolution (at 200 *m/z*). Maven software (Princeton), KEGG pathway database, and an in-house validated standard library (>650 compounds; Sigma Aldrich; IROATech) were used for metabolite assignment and peak integration for relative quantitation. Integrated peak areas were exported into Excel (Microsoft) and elaborated for statistical analysis (*t* test, ANOVA) and hierarchical clustering analysis through the software GraphPad Prism (GraphPad Software) and GENE E (Broad Institute). Top up- and down-regulated metabolic pathways were determined through MetaboAnalyst 3.0 (www.metaboanalyst.ca/MetaboAnalyst/).

### 2.20 Analysis of Bulk RNA Sequence data

For our team’s dataset, the expression values used in these plots correspond to the residuals obtained from a linear regression with gene expression that included all covariates except the disease variable (AAA/control). Gene quantifications were expressed as transcripts per million (TPM), normalized via quantile normalization. Prior to differential expression analysis, genes with low expression levels, defined as TPM values below 0.5 in more than 50% of the samples, were excluded. To identify differentially expressed genes between AAA and controls, a linear regression model was employed, incorporating a number of covariates, including age, sex, flow cell type, flow cell lane, mean GC content, RIN, percentage of RNA fragments > 200 and Qubit. The p-values obtained from the regression were corrected using the Benjamini-Hochberg procedure, and a gene was considered to be differentially expressed if the adjusted p-value was less than 0.05. Samples were clustered into a heatmap by mitophagy-related genes, including “ATG7”, “MAP1LC3B”, “AMBRA1”, “OPTN”, “USP30”, “BNIP3”, “BNIP3L”, “PHB2”, “PINK1”, “FUNDC1”, “ATG5”, “BECN1”, “PARK2”, “PPARGC1A”.

We also analyzed one published AAA dataset, Erik et al. (GEO accession number, GSE 57691), with 20 small and 49 large human AAA samples and 10 donor samples in this cohort. The normalization was applied using limma package. Principal Component Analysis was performed to facilitate clustering of the samples. Outliers were identified and removed. The data matrices were annotated using attached gene symbols. Matrix file was obtained for analyzing Differentially Expressed Genes (DEGs), with |log2FC| > 0.8 and an adjusted p-value < 0.05. Genes were matched with human genes in org.Hs.eg.db. Then GO and KEGG enrichment analysis were conducted. Dot plot and gene-concept network plot were plotted. Finally, samples were clustered into a heatmap by available mitophagy-related genes, including “ATG7”, “MAP1LC3B”, “OPTN”, “USP30”, “BNIP3”, “BNIP3L”, “PHB2”, “FUNDC1”, “ATG5”, “BECN1”, “PPARGC1A”.

### 2.21 Statistics

All results are presented as mean ± standard error of the mean (SEM). Normality was determined using Shapiro-Wilk test. Nonparametric tests were used for data that were not normally distributed. Data with one independent variable were analyzed using Student’s t test or Mann-Whitney U-test. When more independent variables were analyzed, one-way ANOVA or 2-way ANOVA with multiple comparisons were used. Specific statistical tests and *P values* are denoted in figure legends. Two-sided *P values* were used and values <0.05 were considered significant. GraphPad Prism 10.0.0 (GraphPad Software) was used to generate figures and for statistics.

## 3. Results

### mito-QC aged mouse aortas demonstrate decreased mitophagy in VSMCs and endothelial cells and a reduction of PINK1/Parkin expression

Previous studies have demonstrated decreased mitophagy in aged brain and heart tissue, as well as reduced Parkin levels in aged aortas [42–44]. Despite this, changes in mitophagy levels within aged aortic tissue and AAAs have not been explored. We utilized flow cytometry to analyze digested aortas from young (8-10 weeks) and aged (18-20 months) male mito-QC mice using a previously published staining strategy [45] (Fig. 1A, Supplementary Fig. S1A). As gating strategy showed, endothelial cells were identified as CD45^-^CD31^+^ cells, general CD45^+^ cells were identified as CD31^-^CD45^+^ cells, and VSMCs as CD45^-^CD31^-^CD90^-^α-actin^+^ cells (Supplementary Fig. S1A). Plots comparing GFP and mCherry fluorescence across three cellular populations revealed a decrease in GFP levels in CD45^+^ cells from aged aortas and in VSMCs from young aortas. (Fig. 1B). Mitophagy levels were quantified by the mean fluorescence intensity (MFI) of mCherry versus GFP for each group as reported previously by our lab [46]. In endothelial cells, aged mice exhibited a significant decrease in mitophagy of approximately 5%, in both thoracic and abdominal aortic segments compared to young mice (Fig. 1C). Conversely, the CD31^-^CD45^+^ cellular population showed a significant increase in mitophagy levels in aged mice (Fig. 1C). Finally, there was a notable decrease in mitophagy levels of approximately 10%, in the VSMCs of the thoracic and abdominal aged aortic segments (Fig. 1C).

**Figure 1.**
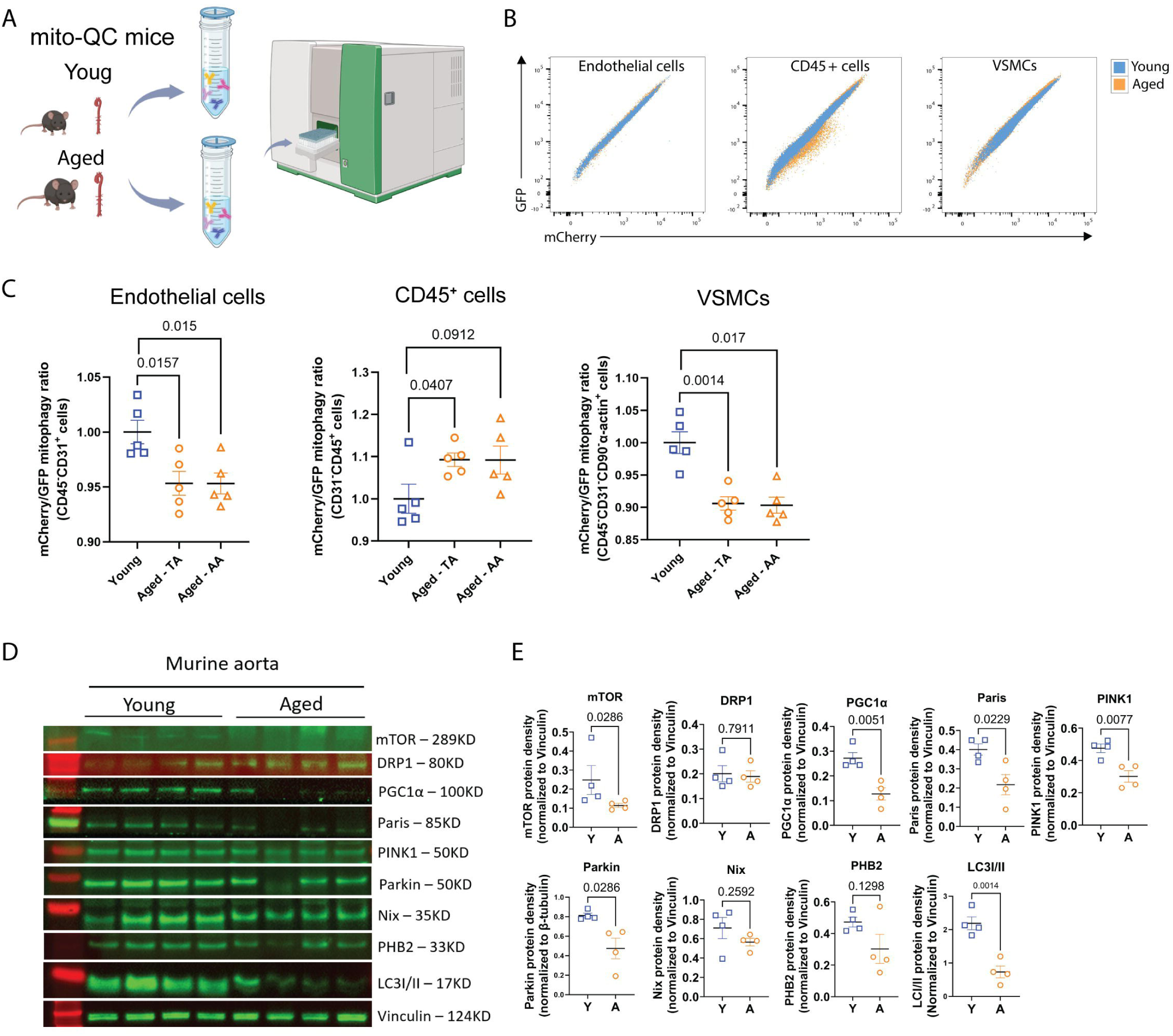
mito-QC aged mouse aortas demonstrate decreased mitophagy in aortic VSMCs and endothelial cells and a reduction of PINK1/Parkin expression. (A) Aortas of young (Y, 8-10 weeks, n=5) and aged (A,18-20 months, n=5) mito-QC male mice were isolated and enzymatically digested into single-cell suspensions, followed by incubation with flow cytometric antibodies conjugated distinct fluorophores. (B) Pseudo color plots for visualizing comparisons of Y and A mitophagy levels (Compensated area for GFP vs. Compensated area for mCherry). (C) Mitophagy levels of different cellular populations between Y and A aorta were quantified by calculating the ratios of mean fluorescence intensity (MFI) of mCherry versus GFP. (D) Protein levels of mTOR, DRP1, PGC1α, Paris, PINK1, Parkin, Nix, PHB2 and LC3I/II in the Y and A aortas were determined by Western Blot (n = 4/group) and normalized to Vinculin (E). Results are presented as mean ± SEM. Unpaired two-tailed Student’s t-test or Mann-Whitney U test were used for statistical analysis.

Western blot analyses of mitophagy-related pathways in the aged aortas demonstrated significant decreases in PINK1 and Parkin expression, whereas changes in Nix and PHB2 were not significant (Fig. 1D, E). Additional mitophagy biogenesis markers, including PGC1α and Paris, and autophagy-related markers LC3II and MTOR, also showed significant reduction via Western blot (Fig. 1D, E). However, the mitochondrial fission marker DRP1 and mitochondrial complexes remained unchanged between young and aged aortas (Fig. D, E, Supplementary Fig. S1B, C). On the other hand, VDAC1, an internal control for mitochondrial complexes, demonstrated a significant decrease in the aged aorta. The specificity of Parkin was further validated in brain and aortic tissues of WT versus Parkin knockout mice (Supplementary Fig. S1D).

### Reduction of Mitophagy in VSMCs in Angiotensin II induced AAAs

We next sought to determine whether AAA induction alters mitophagy using mito-QC male mice in three groups: male mice on a chow diet with saline pump, male mice on HFD with saline pump, and male mice on HFD with AngII infusion at 1000 ng/kg/min. In the HFD/AngII AAA model, serum cholesterol levels gradually increased over a 4-week HFD period (Supplementary Fig. S2A). Mice were harvested 28 days posted AngII infusion, cells were stained as previously mentioned, and flow cytometry was conducted as in Figure 1 (Fig. 2A). The gating strategy was consistent with Figure 1 and adjusted for VSMCs which were categorized into CD45^-^CD31^-^CD90^-^ α-actin^high^ and CD45^-^CD31^-^CD90^-^α-actin^low^ groups, due to altered expression of α-actin in the AAA (Supplementary Fig. S2B-C) [45]. α-actin^high^ and α-actin^low^ VSMCs from the HFD plus AngII group exhibited significantly reduced mitophagy levels compared to the HFD plus saline group (Fig. 2B, C), whereas endothelial cells and CD31^-^CD45^+^ immune cells showed no significant changes (Fig. 2B). In separate studies, mice on an AngII pump without HFD also displayed markedly lower mitophagy in VSMCs compared to Saline pump without HFD (Supplementary Fig. S2D). Additionally, analysis of frozen aortic sections revealed fewer mCherry puncta and reduced colocalization of LAMP-1, a lysosomal marker, and TOMM20, a mitochondrial marker, in the HFD plus AngII group compared to those on HFD plus saline (Fig. 2D, E and F).

**Figure 2.**
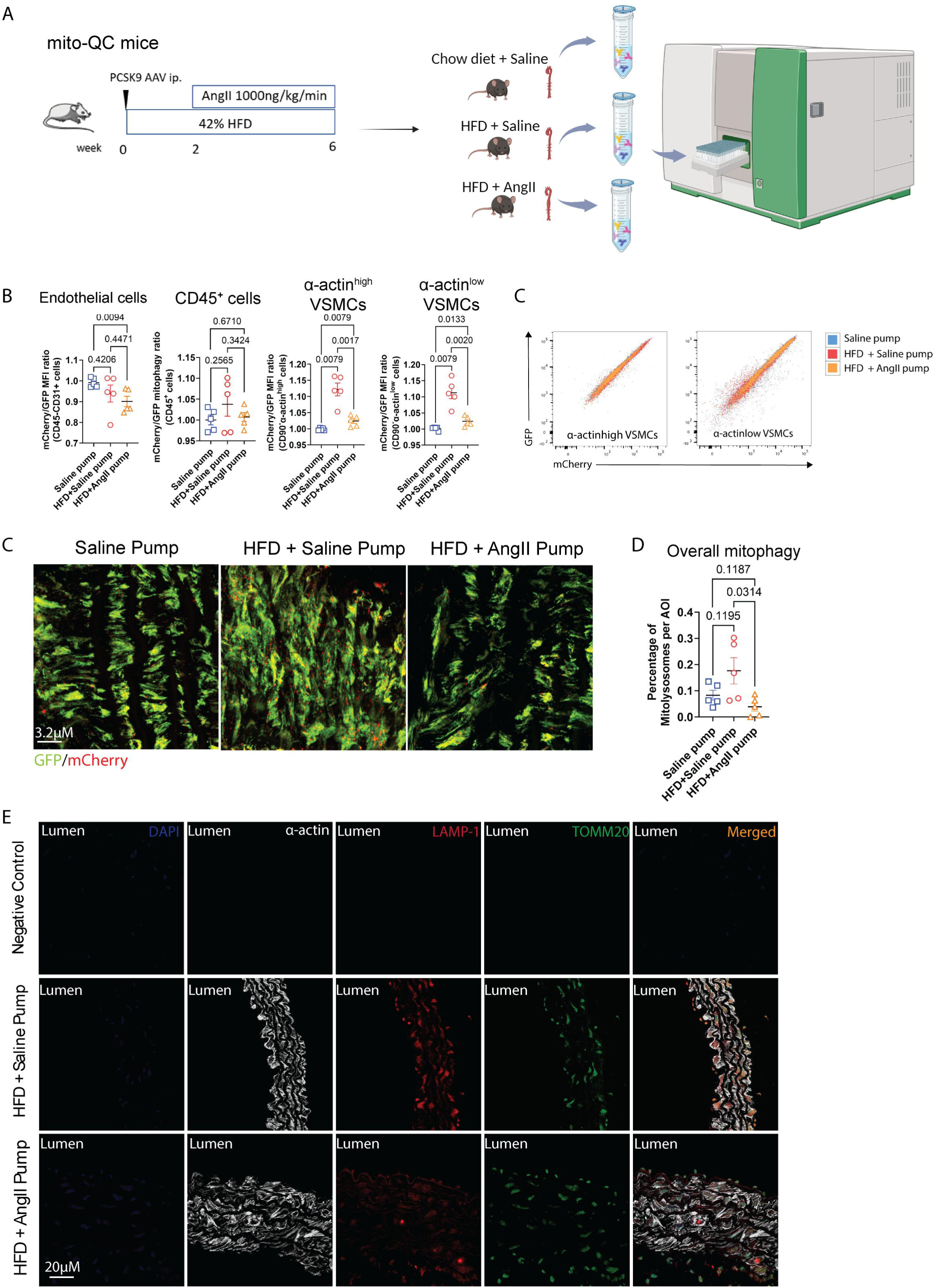
Mitophagy is also Reduced in VSMCs in Angiotensin II induced AAAs. (A) Schematic of the murine AAA AngII model. Briefly, AAV containing murine PCSK9 gain-of function mutation was administrated through a single intraperitoneal injection in conjunction with 42% HFD for 6 weeks. After 2-weeks HFD, the mice were subjected to subcutaneous implantation of an Ang II pump delivering a dosage of 1000ng/kg/min for 4 weeks. Subsequently, aortas from three experimental groups – chow diet with a saline pump, HFD with a saline pump and HFD with an AngII pump (8-12 weeks, n=5 per group, mito-QC mice) – were harvested, digested into single-cell suspensions, and then subjected to the same flow cytometric antibodies as described in Figure 1. (B) Mitophagy levels of endothelial cells (CD45^-^ CD31^+^), CD45^+^ cellular population, α-actin^high^ VSMCs (CD31-CD45-α-actin^high^) and α-actin^low^ VSMCs (CD31-CD45-α-actin^low^) were quantified by calculating the ratios of mean fluorescence intensity (MFI) of mCherry versus GFP. (C) Pseudo color plots for visualizing comparisons of three groups’ mitophagy levels (Compensated area for GFP vs. Compensated area for mCherry). (D) Representative images of frozen-sectioned slides from three groups mentioned above, displaying the GFP and mCherry fluorescence within the aortic tissue. The mCherry puncta represents mito-lysosomes. The quantification of the percentage of mito-lysosomes within ROI is shown in (E). (F) IF staining of α-actin, LAMP-1 and TOMM20 of aortic slides from HFD + Saline or AngII pump group. Results are presented by mean ± SEM. Unpaired two-tailed Student’s t-test or Mann-Whitney U test were used for statistical analysis.

The previous studies demonstrated decreased mitophagy levels in the VSMCs in AAAs following AngII infusion; therefore, we sought to investigate the potential involvement of the PINK1/Parkin-mediated mitophagy pathway in AAAs. qPCR analysis demonstrated Parkin (*Park2*), and α-actin (*ACTA2*) were significantly downregulated in AAA specimens, while *PINK1*, *MYH11*, and other autophagy markers demonstrated no significant changes (Fig. 3A). Western blot analysis noted Parkin and PINK1 were substantially reduced within murine AAAs, whereas PHB2 increased significantly, and Nix remained largely unchanged following HFD and AngII treatment (Fig. 3B, C). Additional Mitochondrial biogenesis and autophagy markers analyzed noted with a reduction in PGC1α, increases in STING, TBK1 and LC3II, alongside a decrease in α-actin in murine AAAs (Fig 3B, C). The downregulation of Parkin and α-actin was further corroborated by IHC staining, with markedly diminished signals in the aortic medial layer (Figure 3D, E and Supplementary Fig. S2C). Furthermore, Western blot analysis confirmed a reduction in SDHB, a complex II protein, in HFD plus Ang II infusion along with a decreasing trend observed for ATP5A in complex V and NDUFB8 in complex I (Fig. 3F, G).

**Figure 3.**
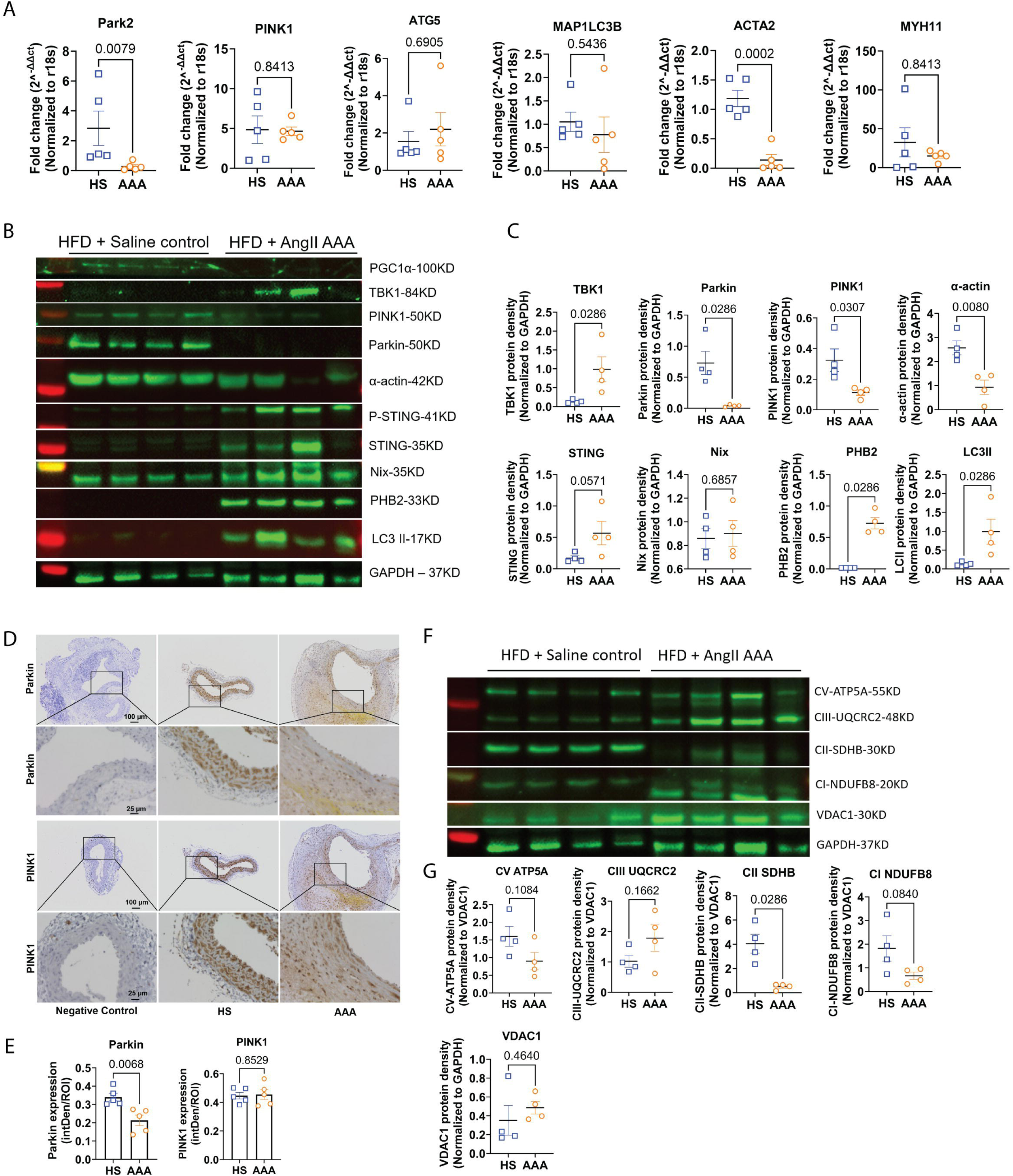
Parkin expression is diminished within the AAA tissue, concomitant with a reduction in the content of mitochondrial complexes. (A) mRNA levels of mitophagy related genes in the aortic tissue of HFD with saline pump (HS) and AAA groups (n=5 per group), including Park2, ACTA2, MYH11, ATG5, MAP1LC3b and PINK1, with r18s as housekeeping gene. (B-C) Protein levels in the aortic tissue of HS and AAA groups, including PGC1α, TBK1, PINK1, Parkin, α-actin, STING, Nix, PHB2 and LC3II were determined by Western blot (n=4 per group), employing GAPDH as internal control for quantification. (D-E) Representative aortic IHC staining images and quantification of Parkin and PINK1 in the HS and AAA group (n=5 per group). (F-G) The contents of four representative proteins associated with mitochondrial complexes within the aortic tissues in HS and AAA group, including complex V - ATP5A, complex III - UQCRC2, complex II - SDHB, complex I - NDUFB8, were determined by Western blot, utilizing VDAC1 as internal control for quantification. Results are presented by mean ± SEM. Unpaired two-tailed Student’s t-test or Mann-Whitney U test were used for statistical analysis.

### Human AAA samples exhibit decreased expression of mitophagy mediator Parkin and PINK1 in gene and protein level

The expression of mitophagy-related genes was analyzed in our bulk RNA-seq dataset, which included 44 control samples and 96 AAA samples (Fig. 4A). PINK1 exhibited a significant decrease in the AAA group and PARK2 showed a decreasing trend. Similarly, USP30, an antagonist of the PINK1/Parkin mitophagy pathway, also exhibited a decreasing trend. Mediators of other mitophagy pathways, such as BNIP3 and BNIP3L, were downregulated, but PHB2 expression remained unchanged. For generic autophagy mediators, MAP1LC3B and ATG7 were upregulated, whereas ATG5 showed no significant change. Regulators of the mitophagy pathway, including AMBRA1, BECN1, FUNDC1, and OPTN, were decreased in the AAA group. The mitochondrial pathogenesis-related gene, PPARGC1A, showed no change.

**Figure 4.**
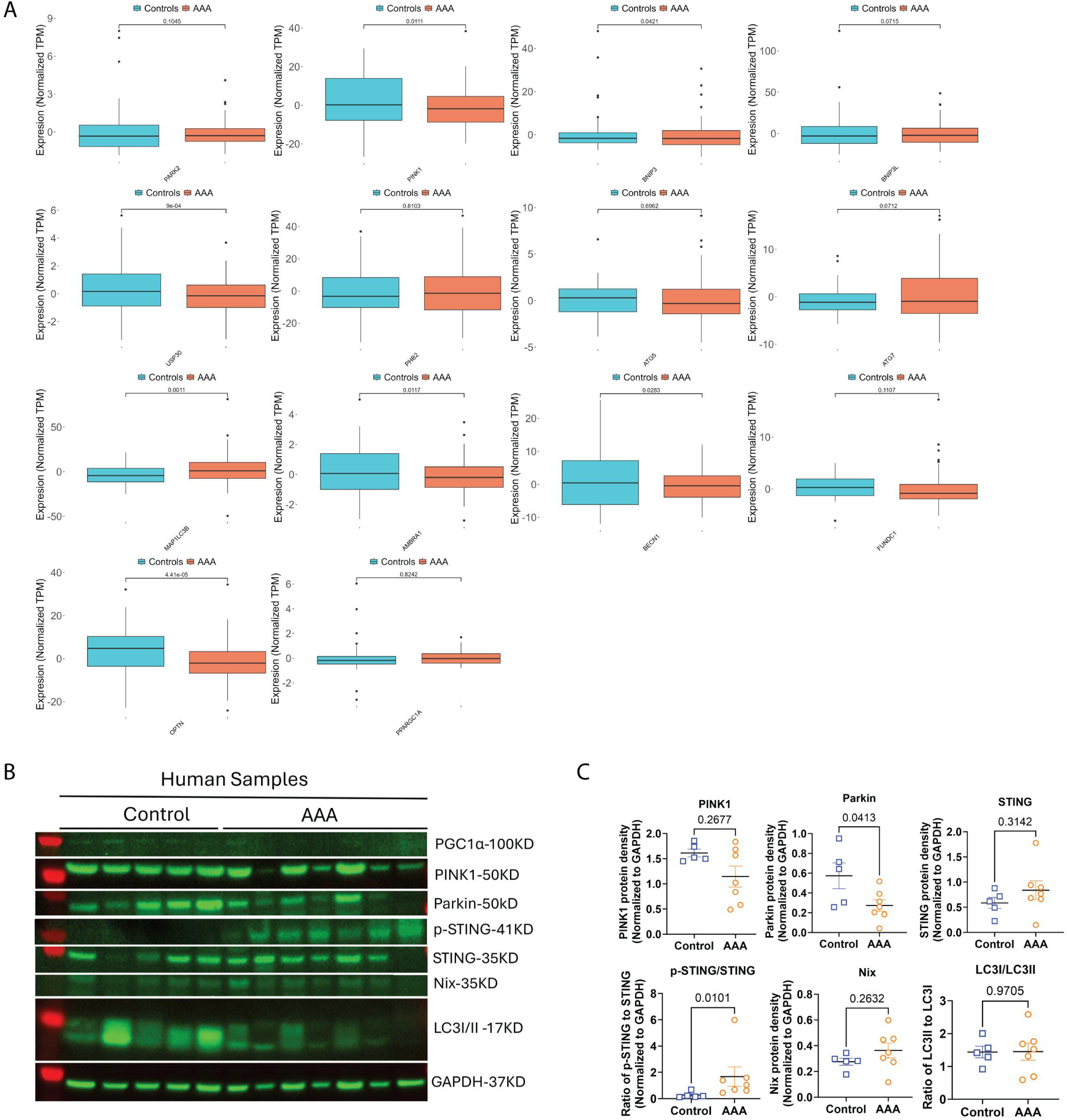
Mitophagy mediators PINK1 and Parkin exhibited a decreasing trend in both gene and protein expression levels in human AAA tissues. (A) Gene expression analysis of mitophagy-related genes (including PARK2, PINK1, BNIP3, BNIP3L, USP30, PHB2, ATG5, ATG7, MAPLC3B, AMBRA1, BECN1, FUNDC1, OPTN, and PPARGC1A) was conducted on human control (n=44) and AAA (n=96) tissues using bulk RNA sequencing. (B-C) Protein levels of PGC1α, PINK1, Parkin, Sting, Nix, and LC3I/II were assessed in human aortic tissue from healthy aorta and AAA groups (n=5-7 per group) by Western blot, with GAPDH used as an internal control for quantification. Results are presented as mean ± SEM. Statistical analysis was performed using an unpaired two-tailed Student’s t-test or Mann-Whitney U test.

In this dataset, the control and AAA groups could not be distinctly clustered based on these mitophagy-related genes (Supplementary Fig. S3A). In the publicly available dataset GSE57691, a clearer distinction between the control and AAA groups was observed using available mitophagy-associated genes (Supplementary Fig. S3B). KEGG pathway enrichment analysis (Supplementary Fig. S3C) indicated that genes involved in biological processes such as aerobic respiration, the electron transport chain, oxidative phosphorylation, ATP synthesis coupled with electron transport, and the mitochondrial ATP synthesis coupled electron transport and aerobic electron transport chain were downregulated in the AAA group. Additionally, genes associated with cellular components, including the respirasome, respiratory chain complex, and mitochondrial respirasome, were downregulated. Genes related to molecular functions, such as electron transfer activity and antioxidant activity, were also downregulated.

A gene-concepts plot revealed that the observed decreases in ATP synthesis coupled with electron transport and the electron transport chain were primarily due to downregulated proteins in the mitochondrial complex, which were also associated with the response to peptide hormones via gene CYC1 (Supplementary Fig. S3D). In our human bulk RNA-seq dataset, a significant downregulation of nuclear genes encoding mitochondrial complexes I-V (Supplementary Fig. S3E-I) and mitochondrial ribosome components (Supplementary Fig. S3J-K) was observed in the AAA samples. Protein expression analysis demonstrated that Parkin was significantly decreased in the AAA samples, with PINK1 also showing a decreasing trend. There were no significant changes in STING, Nix, and the LC3II/I ratio (Fig. 4B-C).

### Parkin SMCKO mice have increased propensity for AAA rupture and an augmented maximal diameter of the aneurysm, concomitant with enhanced mitochondrial dysfunction

Given reduced mitophagy levels in VSMCs in AAAs and reduced Parkin expression in the aortic medial layer following AngII infusion, we next sought to examine effects of cell specific Parkin elimination in VSMCs in AngII AAAs. We utilized *Myh11-creER^T2^Parkin^fl/fl^* in an HFD/AngII-induced AAA model [28, 30], by administering either vehicle (Control) or tamoxifen (Parkin SMCKO) followed by AAA induction (Fig. 5A). Verification of Parkin knockout in VSMCs was confirmed through genotyping of tail samples and Western blot analysis of the mouse aorta (Supplementary Fig. S4A, B). Prior to AngII pump implantation, the Parkin SMCKO group exhibited significantly lower serum cholesterol levels and weight gain compared to the control group; however, no differences in cholesterol or weight gain were observed at the time of harvest 6 weeks following tamoxifen administration (Supplementary Fig. S4C, D). Following AngII infusion, aortic ruptures were confirmed by autopsy and were included as a death event in the survival analysis. The Parkin SMCKO group exhibited lower survival rates (7/39 Parkin SMCKO versus 1/35 WT; p=0.0383), a trend toward greater aneurysm incidence (p=0.066), and increased maximal aortic diameters compared to WT controls (Fig. 5B, C). In the elastase-induced AAA model, increased aortic dilation was also observed in the Parkin SMCKO group compared to the control group, with mean dilation percentages of 136.80 ± 20.80% versus 99.69 ± 22.33% (p=0.0009; Supplementary Fig. S4E).

**Figure 5.**
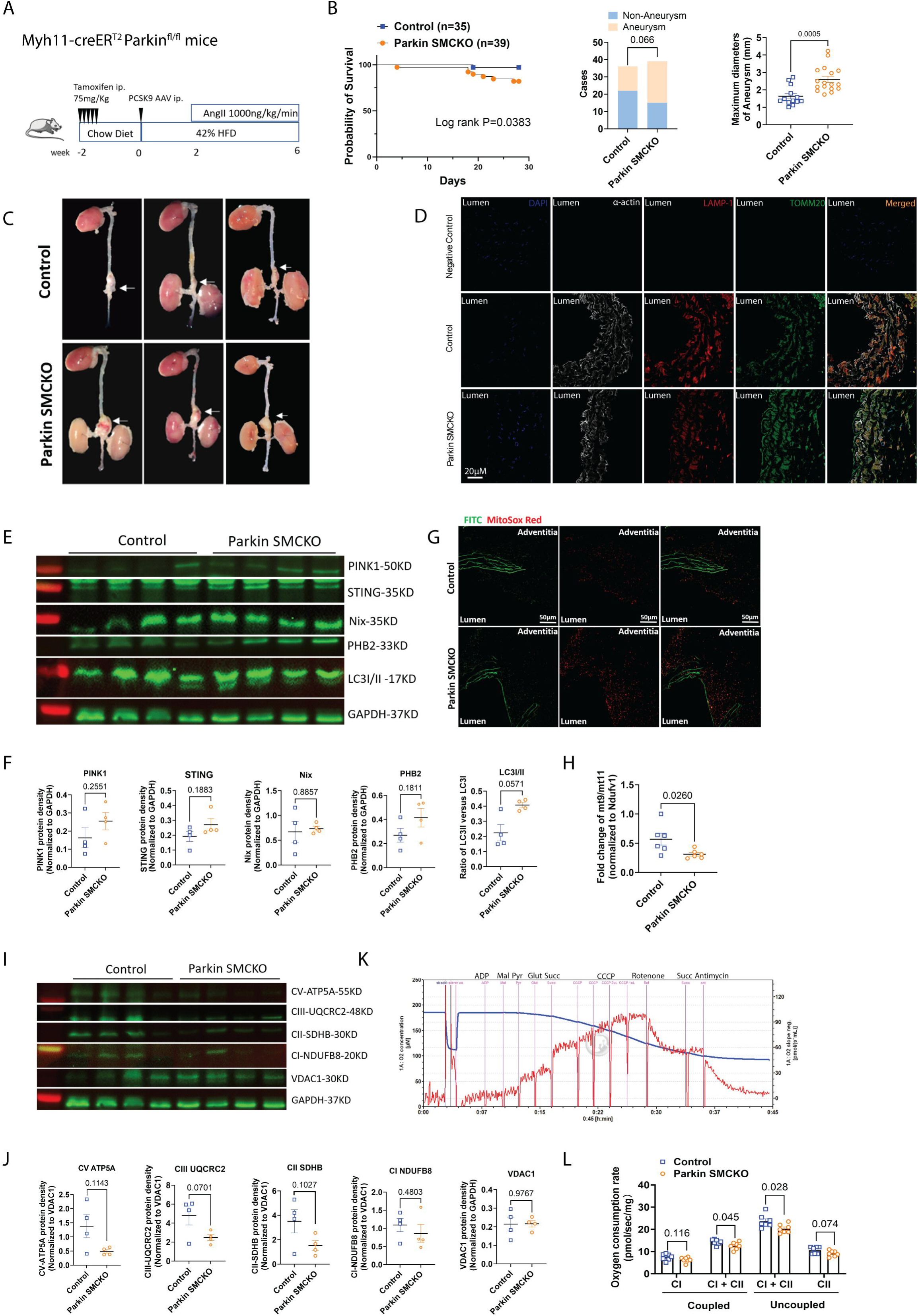
Parkin deficiency in the VSMCs is associated with an increased propensity for AAA rupture and augmented maximal aortic diameter, concomitant with enhanced mitochondrial dysfunction. (A) Schematic of AngII AAA model in Myh11-creERT2 Parkin fl/fl mice cohort. Tamoxifen was administered via intraperitoneal injection at a daily dosage of 75mg/kg for 5 days. After a 9-days interval, the standard procedure for AAA was implemented. (B) Kaplan-Meier survival curves, AAA incidence rate, and maximal aneurysmal diameters in control (n=35) versus Parkin SMCKO (n=39) groups. Log rank test was used for survival analysis. Fisher’s exact test was used for incidence analysis. (C) Representative images of dissected aortas were presented from control and Parkin SMCKO groups. (D) IF staining of α-actin, LAMP-1 and TOMM20 on aortic slides from control vs. Parkin SMCKO group. (E-F) Protein levels in the aneurysms of control and Parkin SMCKO groups, including PINK1, STING, Nix, PHB2, LC3II/I and GAPDH, were determined by Western blot (n=4 per group), employing GAPDH as internal control for quantification. (G) Representative images of frozen-sectioned AAA slides stained with MitoSoxred from control vs. Parkin SMCKO groups are displayed. (H) mtDNA copy numbers in the AAAs of control vs Parkin SMCKO groups were determined by evaluating the fold change ratios of mt9/mt11 versus Ndufv1 respectively, a gene in mtDNA and nuclear DNA encoding the proteins for mitochondrial complex I. (I-J) The contents of four representative proteins associated with mitochondrial complexes within the aortic tissues in control and Parkin SMCKO group, including complex V - ATP5A, complex III - UQCRC2, complex II - SDHB, complex I - NDUFB8, were determined by Western blot, utilizing VDAC1 as internal control for quantification. (K) Representative protocol for measuring the maximal oxygen consumption rate (OCR) using substrates, inhibitors and uncoupler (see methods). (L) Maximal aortic OCR with substrates for complex I and I+II coupled oxidative phosphorylation (OXPHOS) and maximal OCR for complex I+II and complex II in uncoupled conditions, using the AAA tissues from control and Parkin SMCKO groups. 2-way ANOVA with Sidak’s multiple comparison test were used for **L**. Results are presented by mean ± SEM. Unpaired two-tailed Student’s t-test or Mann-Whitney U test were used for statistical analysis for **B (3)**, **I**, **F** and **J**. Results are presented by mean ± SEM. Unpaired two-tailed Student’s t-test or Mann-Whitney U test were used for statistical analysis for **B (3)**, **I**, **F** and **J**.

At the tissue level, less colocalization of LAMP1 and TOMM20 was observed in the Parkin SMCKO group (Fig. 5D). Western blot analysis revealed an increase in STING protein levels and an altered autophagy marker ratio of LC3II to LC3I in the Parkin SMCKO group (Fig. 5E, F). Furthermore, Parkin SMCKO exhibited increased mitochondrial ROS, decreased mitochondrial copy numbers (p=0.0260), and a trending decline in protein levels of ATP5A in complex V (p=0.140), UQCRC2 in complex III (p=0.0701), and SDHB in complex II (p=0.1027; Fig. 5I-J). The protocol employed for measuring oxidative phosphorylation (OXPHOS) capability in aortic mitochondria is detailed schematically in Figure 5K [39], and demonstrated that the oxygen consumption rate, both coupled and uncoupled for Complex I and II, was significantly reduced in the Parkin SMCKO group compared to controls (Fig. 5L).

### Parkin SMCKO AAA mice exhibited metabolomic profiles with altered ATP synthesis, nucleotide metabolism, increased oxidative stress, deficient fatty acid β-oxidation

Serum metabolomics analysis of control versus Parkin SMCKO groups following AngII AAA formation was performed, and bioinformatics analyses was conducted. The scores plot (Fig. 6A) reveals a clear separation between the two groups, indicating significant metabolic differences due to Parkin SMCKO. The heatmap, volcano plot, and analysis of individual metabolites (Fig. 6B, C and Supplementary Fig. S4F) show notable increases in phosphate and diphosphate levels, suggesting alterations in ATP synthesis. Additionally, levels of UDP and xanthine were also elevated, reflecting changes in nucleotide metabolism and increased oxidative stress. Elevated acyl-C4-DC levels point to deficient fatty acid β-oxidation, while increased spermidine indicates a compensatory response in autophagy. Metabolomic pathway analysis (Fig. 4P) also highlights significant changes in metabolites involved in both glycolysis and the TCA cycle, including 2/3-phospho-D-glycerate, 1,3-bisphospho-D-glycerate, and oxaloacetate.

**Figure 6.**
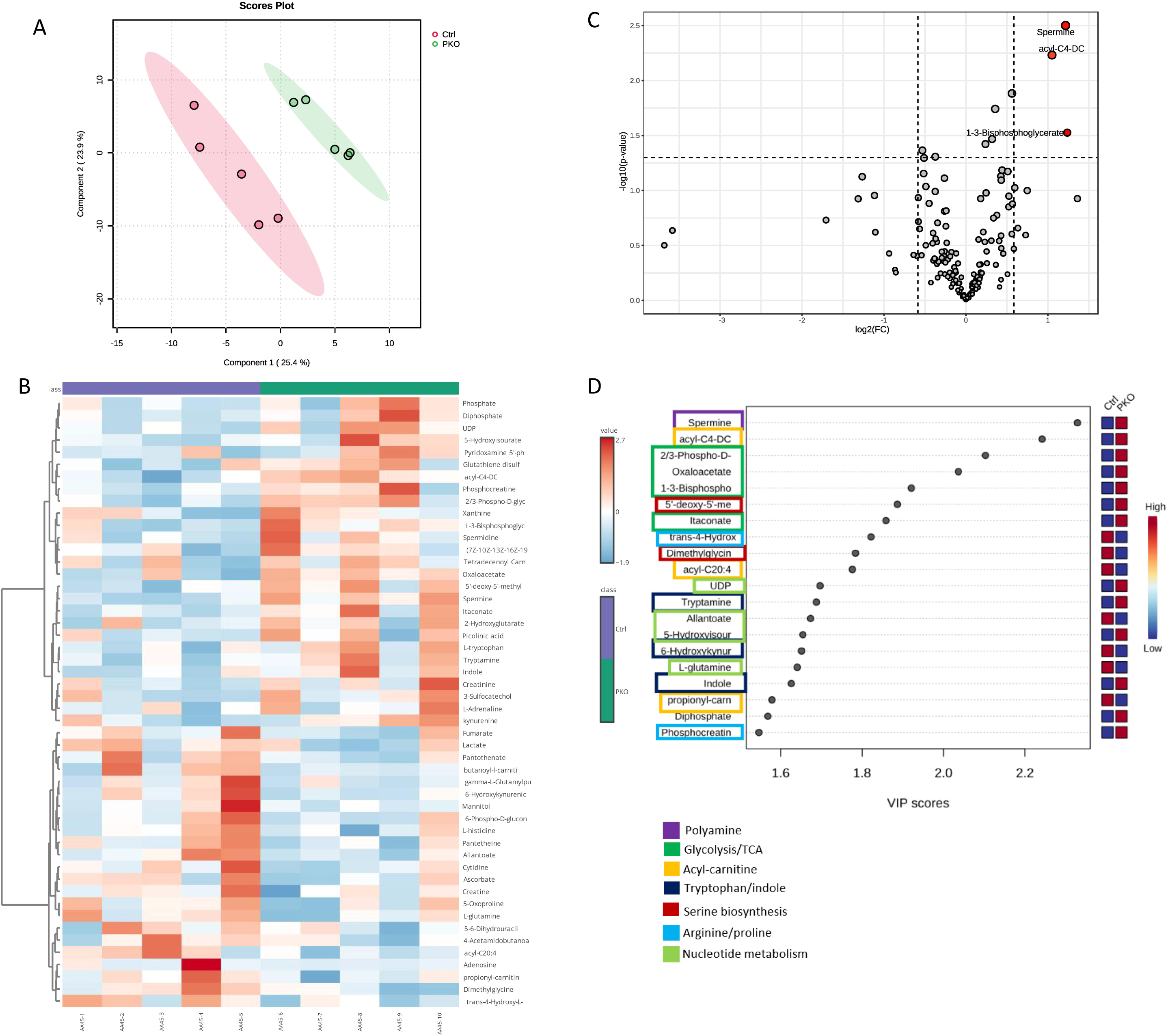
Serum metabolomics data showed mitochondrial dysfunction in Parkin SMCKO AAA mice. Metabolomic analysis was performed on serum samples from ctrl and Parkin SMCKO group (n=5 per group). PLS-DA analysis was performed after normalization of the values (A). The heat map was plotted by top 50 metabolites from two groups (B). The Volcano map was plotted with the threshold as p-value≤0.05 and Log2FC≥ ± 1.5 (C). VIP plot showed the scores of VIP features in different metabolomic pathways that discriminate between Control and Parkin SMCKO group (D).

### VB-08, a USP30 inhibitor, can improve the reduction in mitophagy, and mitochondrial dysfunction caused by AngII in vitro, with effect dependent on Parkin

Primary VSMCs isolated from the aortas of mito-QC mice were treated with AngII at varying concentrations with the proportion of lysosomal mitochondria, indicated by diminished percentage of GFP signal, demonstrating a dose-dependent decrease following AngII treatment (Fig. 7A). In separate studies, MOVAS cells treated with AngII exhibited an increase in mitochondrial ROS that was not dose-dependent (Fig. 7B). In subsequent studies, when MOVAS cells were co-treated with AngII and VB-08, a highly selective USP30 inhibitor small molecule and enhancer of the PINK1/Parkin mitophagy pathway, the AngII-induced reduction in mitophagy was significantly reversed (p=0.0004 AngII vs. AngII + VB-08; Fig. 7C). The co-treatment led to a trend toward a reduction in mitochondrial ROS (p=0.0770; Fig. 7D). Concurrently, the MOVAS cells displayed a decrease in total mitochondrial content but an increase in mitochondrial health, as evidenced by the ratio of mitochondrial membrane potential (TMRM) to Mitotracker fluorescence (Fig. 7E). Western blot analyses demonstrated that treatment with AngII notably decreased Parkin expression, particularly at concentrations of 100 nM and 500 nM (Fig. 7F, supplementary Fig. S5A, B). Conversely, treatment with VB-08 significantly increased the expression of both PGC1α and Parkin (Fig. 7G). Protein expression analysis demonstrated that Parkin was significantly decreased in the AAA samples, with PINK1 also showing a decreasing trend. To demonstrate whether the protective effect of VB-08 is related to Parkin expression, we used lentiviral Parkin shRNA to knock down Parkin protein in MOVAS by around 43% (supplementary Fig. S5C, D). The cells were treated with AngII alone or in combination with VB-08. In control (scrambled shRNA) and Parkin knockdown MOVAS, AngII increased mitochondrial ROS levels and reduced the percentage of healthy mitochondria. While VB-08 was expected to reduce ROS and promote the mitochondrial health, it left notably higher ROS levels and lower percentage of healthier mitochondria in the Parkin knockdown MOVAS (Fig. 7H).

**Figure 7.**
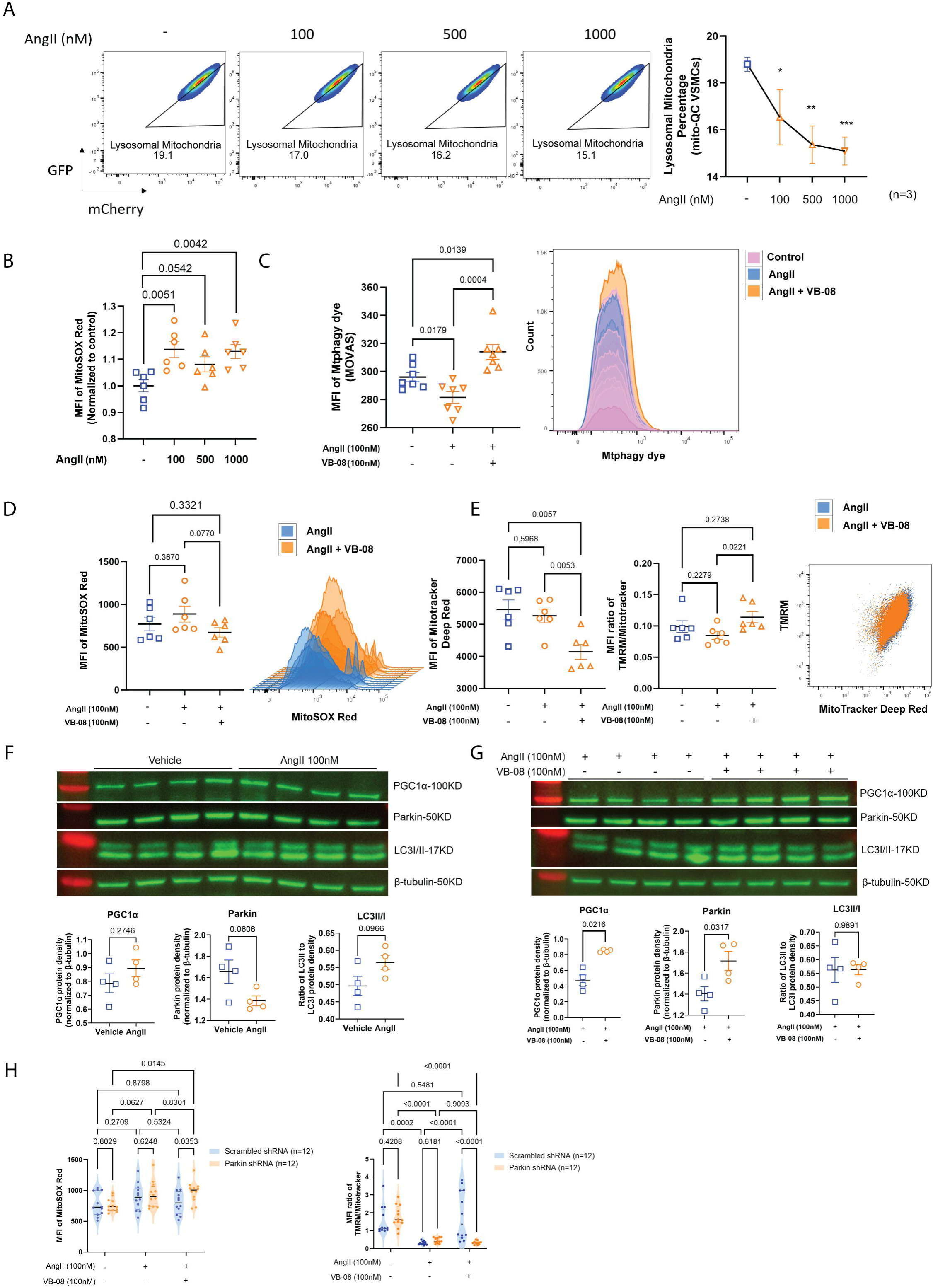
Angiotensin II reduces mitophagy levels of VSMCs, leading to increased mtROS and mitochondria dysfunction, while USP30 inhibition via VB-08 ameliorates the outcome in vitro. (A) Pseudocolor plots for visualizing the mitophagy levels by gating the cells with lower MFI ratio of GFP versus mCherry. Cells are primary VSMCs isolated from Mito-QC mice, treated with 100nM, 500nM and 1000nM AngII (n=3 per group). (B) The MFI of MitoSox Red in MOVAS treated with 100nM, 500nM, and 1000nM AngII (n=6 per group). (C-E) The MFI of Mtphagy dye (n=7 per group) and MitoSox Red (n=6 per group) within MOVAS respectively treated with vehicle, 100nM AngII, or a combination of 100nM AngII with 100nM VB-08, were quantified and showed in a histogram, same for the MFI of Mitotracker and TMRM. The MFI ratio of TMRM to Mitotracker was employed as a metric for assessing the proportion of healthy mitochondria within the cells. (F-G) Alteration of protein levels in MOVAS after treatment with AngII alone or AngII in combination with VB-08, including PGC1α, Parkin, LC3I/II, were determined by Western blot (n=4 per group). β-tubulin is the internal control for normalization. (H) For MOVAS transfected with lentiviral scrambled or Parkin shRNA, the MFI of MitoSox Red within cells treated with vehicle, 100nM AngII, or a combination of 100nM AngII with 100nM VB-08 (n=12 per group), were quantified, same for the MFI ratio of TMRM to Mitotracker. Results are presented as mean ± SEM. Unpaired two-tailed Student’s t-test or Mann-Whitney U test were used for statistical analysis. Two-way ANOVA was used for **H**.

### Treatment with USP30 inhibitor VB-08 in vivo significantly reduced aneurysm incidence and diameter and enhanced mitochondrial function

mito-QC mice, subjected to a HFD and AngII-induced AAA model, received either vehicle or VB-08 treatment daily, starting five days before and continuing four weeks after AngII pump implantation (Supplementary Fig. S6A). Subsequent flow cytometry analysis of harvested aortas revealed an increase in mitophagy levels of VSMCs with VB-08 treatment (Supplementary Fig. S6B). This regimen was replicated in C57BL/6 mice with the AAA model over the same duration (Fig. 8A). Treatment with VB-08 and AngII infusion demonstrated no significant differences in serum cholesterol levels between the two groups (Supplementary Fig. S6C) nor were there survival differences (Fig. 8B). However, AAA incidence was significantly lower in the USP30 inhibitor group compared to vehicle control (p=0.010; Fig. 8B) with a corresponding trend toward decreased aneurysm diameter (p=0.0652; Fig. 8B, C). Protein expression analysis via Western blot in the aorta indicated a significant increase in PINK1 levels (p=0.0164) and the ratio of LC3II to LC3I (p=0.0011) in the USP30 inhibitor group (Fig. 8D, E). While STING expression decreased significantly (p=0.0208), Parkin expression exhibited a downward trend (p=0.0626; Fig. 8D, E). Conversely, PGC1α levels trended upward (p=0.2976), with no notable changes in Nix and PHB2 expressions (Fig. 8D, E). mtDNA copy numbers in the aorta markedly increased in the USP30 inhibitor group (p=0.0426; Fig. 8F). Following the established protocol described for Parkin SMCKO, both uncoupled and coupled oxygen consumption rates in complex I and II showed an upward trend in the USP30 inhibitor-treated group, although no significant changes were observed in complex content except that ATP5A in complex V showing an increasing trend (Figure 8G, supplementary Fig. S6D, E).

**Figure 8.**
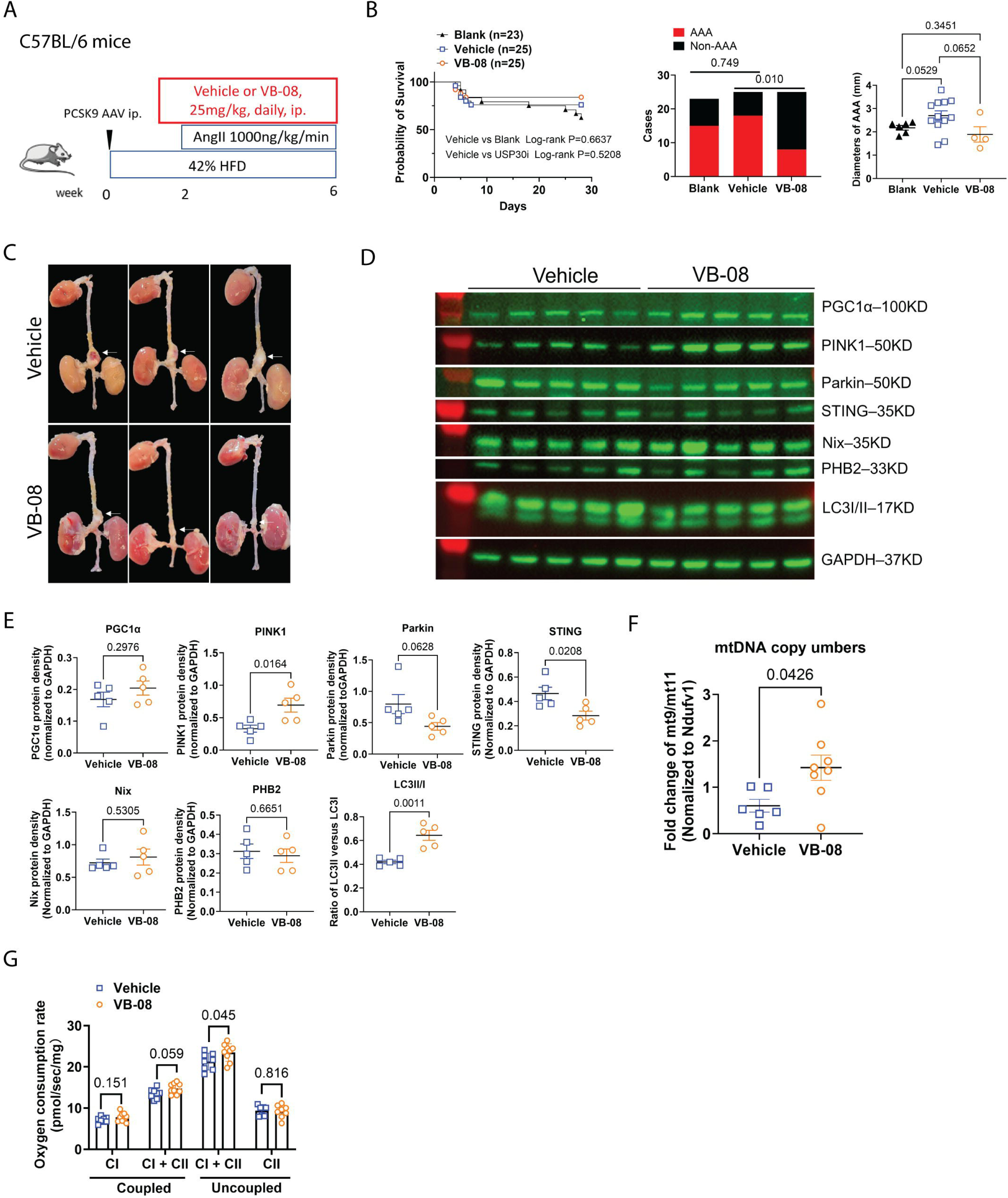
VB-08, a USP30 inhibitor, reduces aneurysm incidence and maximal aortic diameters in vivo during Ang II infusion. (A) The strategy for inducing and therapeutically targeting AAA with VB-08 in C57BL/6 male mice cohort. (B) Kaplan-Meier survival curve, AAA incidence rate, and maximal diameters of aneurysms in blank (n=23), vehicle (n=25), and VB-08 (n=25) groups. Log rank test was used for survival analysis. Fisher’s exact test was used for incidence rate analysis. (C) The representative images of dissected aortas in the vehicle vs. VB-08 group. (D-E) Protein levels in the descending aortas of vehicle vs. VB-08 inhibitor groups, including PGC1α, PINK1, Parkin, STING, Nix, PHB2, LC3II/I and GAPDH, were determined by Western blot (n=5 per group), employing GAPDH as internal control for quantification. (F) mtDNA copy numbers in the aorta of vehicle vs. VB-08 treated group were determined by evaluating the fold change ratios of mt9/mt11 versus Ndufv1. (G) Maximal aortic OCR with substrates for complex I and I+II coupled oxidative phosphorylation (OXPHOS) and maximal OCR for complex I+II and complex II in uncoupled conditions, using the aortic tissues from vehicle vs. VB-08 groups (n = 8-9 per group).

### AAAs with USP30 inhibition exhibited metabolomics profile with altered ATP synthesis, energy metabolism, neurotransmitter turnover, nucleotide metabolism, and decreased oxidative stress

Serum metabolomics analysis of the vehicle (control) and VB-08 groups, as demonstrated by the scores plot (Fig. 9A), shows a clear separation between groups, indicating significant metabolic differences due to VB-08 treatment. Metabolomic profiling, as depicted in the heatmap, volcano plot, and individual metabolite analysis (Fig. 9B, C, and Supplementary Fig. S6F), reveals distinct metabolic shifts. Decreased levels of phosphate and acyl-carnitine, which are involved in ATP synthesis and fatty acid oxidation [47, 48], contrast with the trends observed in the Parkin SMCKO intervention, highlighting a unique and connected metabolic response to USP30 inhibition and Parkin SMCKO. Notably, pyruvate and 2-oxoglutarate levels are reduced, while fumarate is increased, indicating a shift in energy metabolism from glycolysis and the citric acid cycle towards oxidative phosphorylation with VB-08 administration. Changes in lipid and carbohydrate metabolism are suggested from decreased hexanoic acid and D-rhamnose, alongside increased gamma-L-glutamyl-D-glucose and N-acetylneuraminic acid, reflecting adjustments in fatty acid oxidation and glycosylation [49, 50]. Alterations in nucleotide and amino acid metabolism are marked by decreased cytidine and hypoxanthine levels, coupled with increased GMP and L-methionine, suggesting enhanced nucleotide synthesis, RNA production, and DNA methylation (supplementary Fig. S6F) [51, 52]. Additionally, there are notable changes in neurotransmitter-related metabolites, with decreased acetylcholine versus increased 5-hydroxyindoleacetate and serotonin, indicating altered cholinergic and serotonergic dynamics. Elevated levels of S-glutathionyl-L-cysteine, ascorbate, and Cys-Gly highlight an up-regulation in antioxidant defenses and detoxification processes, reducing oxidative damage and supporting cellular health (supplementary Fig. S6F) [53]. These findings are further corroborated by the metabolomic pathway analysis (Fig. 9D), which underscores significant alterations across various metabolic pathways, illustrating a broad metabolic reprogramming driven by USP30 inhibition.

**Figure 9.**
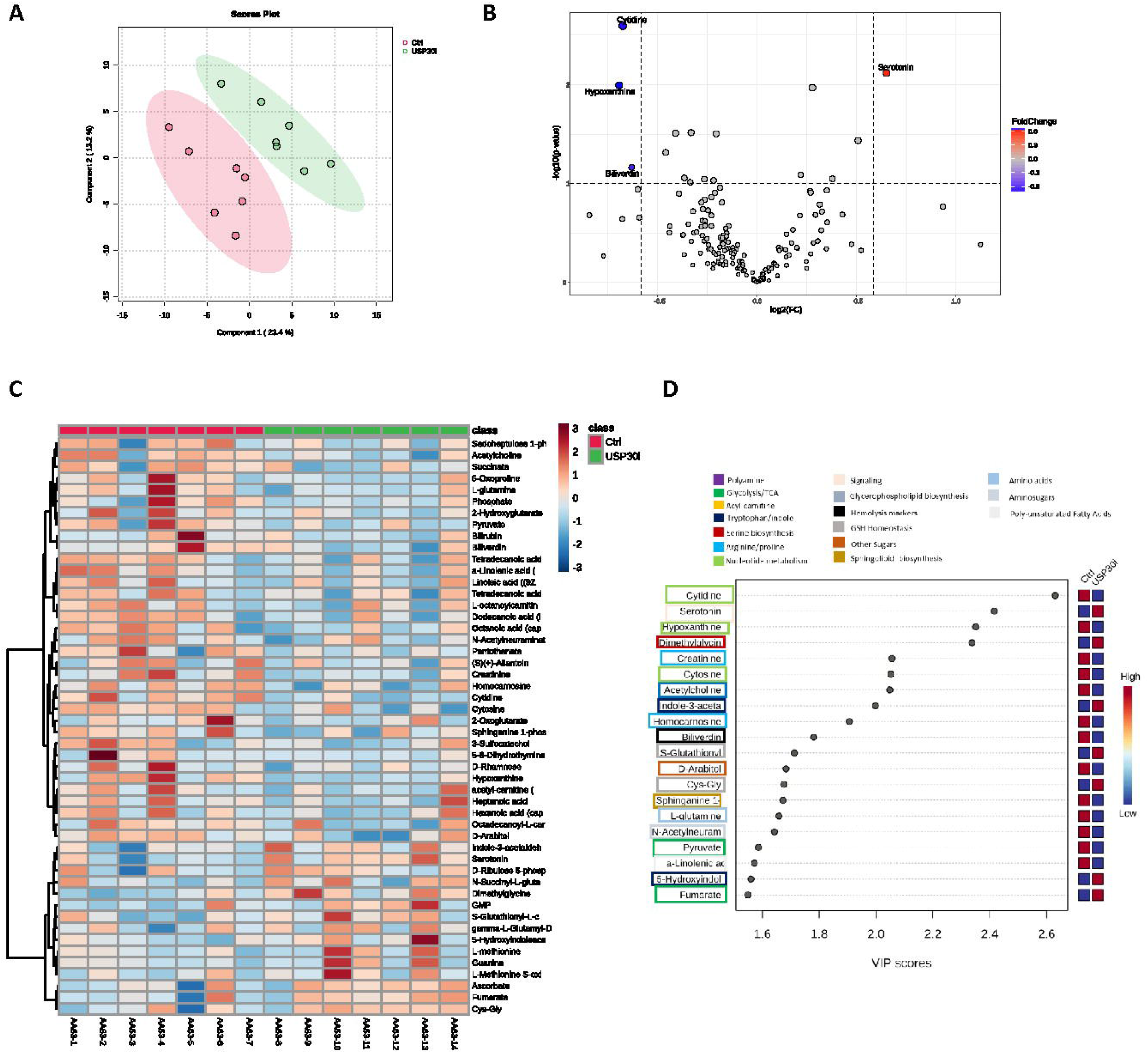
Serum metabolomics data showed altered metabolites in AAA mice following USP30 inhibition by VB-08. Metabolomic analysis was performed on serum samples from vehicle and VB-08 group (n=7 per group) in AAA mice. PLS-DA analysis was performed after normalization of the values (A). The Volcano map was plotted with the threshold as p-value≤0.05 and Log2FC≥ ± 1.5 (B). The heat map was plotted by top 50 metabolites from two groups (C). VIP plot showed the scores of VIP features in different metabolomic pathways that discriminate between vehicle and VB-08 group (D).

## 4. Discussion

Recent evidence suggests that the dysregulation of mitochondrial quality control during aging adversely affects mitochondrial function and may contribute to cardiovascular disease progression [54, 55]. Evidence also suggests that mitophagy acts as a molecular sensor for the maintenance of mitochondrial quality by clearing defective mitochondria. Although evidence suggests decline in mitophagy levels is a hallmark of aging, alterations in mitophagy within specific aortic cell types have not been reported. Our study shows that mitophagy declines in aged murine aortas specifically in VSMCs (CD90^-^α-actin^+^) and endothelial cells (CD31^+^), but not in general immune cells (CD45^+^). Furthermore, while it is known that the prevalence and severity of AAA increases with advancing age, few studies have investigated Parkin-dependent mitophagy in AAAs [15, 19]. Our findings reveal that mitophagy decline in VSMCs during AAA development is associated with reduced expression of PINK1/Parkin-mediated mitophagy pathway in the aorta, a known mitophagy regulatory pathway, in both murine and human AAAs. We demonstrated Parkin SMCKO exacerbates murine AngII and elastase induced AAAs, leading to increased severity of mitochondrial dysfunction and STING expression, underscoring the importance of Parkin-mediated mitophagy in AAA. In contrast, enhancing the PINK1/Parkin-mediated mitophagy pathway through USP30 inhibition significantly reduces AAA incidence, improves mitochondrial function, and decreases STING expression, indicating that the PINK1/Parkin mitophagy pathway is a promising therapeutic target for AAA. Our data suggest that enhancing the PINK1/Parkin-mediated mitophagy pathway in VSMCs can improve AAA outcomes, positioning it as a potential therapeutic avenue for this aging-related disease.

We initially identified decreased mitophagy levels in the VSMCs and endothelial cells of aged murine aortas (20-month-old mito-QC mice), while general immune cells demonstrated increased mitophagy levels. This increase in general immune cell mitophagy is logical considering that senescent immune cells in aged aortas are more inflammatory, relying on the turnover of defective mitochondria to facilitate inflammatory responses [56, 57]. Previous research using mt-Keima mice reported increased overall aortic mitophagy in 16-month-old male mice, although failed to distinguish cellular types [39]. Both mito-QC and mt-Keima mice are mitophagy reporter tools that utilize pH-sensitive fluorescence detection. The fixable characteristic of mito-QC mice, unlike the live-imaging feature of mt-Keima mice, gives mito-QC method the ability to measure mitophagy levels in different cellular types following fixation. Previous studies have demonstrated good coordination between these two tools for mitophagy measurements[58, 59]. Moreover, mito-QC mice exhibit longer lifespans than mt-Keima mice, making them more applicable for *in vivo* aging studies. Consistent with our results, Thomas et al. found reduced Parkin expression in the aortas of 27-28-month-old mice, attributing this impairment to increased mitochondrial redox stress and superoxide, which is also implicated in AAA pathogenesis [42, 60, 61].

Mitophagy in VSMCs is suggested to protect cells from apoptosis and oxidative stress caused by cytochrome C release and damaged mitochondria [62]. Despite the verified importance of autophagy in maintaining VSMC homeostasis in AAA, verified by Atg5 or Atg7 SMCKO exacerbating AAA, little is known about the role of VSMC mitophagy during AAA development [45, 63]. Our study demonstrates that mitophagy decline in VSMCs occurs during AAA development and removal of Parkin within VSMCs increases AAA incidence. Furthermore, the impaired mitochondrial respiratory function, decreased healthy mitochondria proportion, and increased mitochondrial reactive oxygen species (mtROS) in Parkin SMCKO AAA underscore Parkin’s importance in AAA. Our current findings corroborate previous studies that demonstrated decreased Parkin expression in human AAA samples compared to healthy controls [64].

The cGAS-STING pathway, activated by cytosolic DNA, including leaked mtDNA segments, is known to partially prevent AAA development when inhibited [13]. Defective mitophagy during aging has been shown to activate the mtDNA-related cGAS-STING pathway[17, 65, 66]. Our results show decreased PINK1 and Parkin expression alongside increased STING in AAA samples. In Parkin SMCKO AAA, where mitochondrial dysfunction is more severe, STING expression is further elevated, while enhancing PINK1/Parkin-mediated mitophagy through USP30 inhibition decreases STING expression. This highlights a link between PINK1/Parkin-mediated mitophagy and the STING-related innate immune pathway in AAA, providing additional evidence for the investigation of mitophagy, aging, and STING interactions.

There is evidence suggesting crosstalk between different mitophagy mechanisms, although the interplay remains to be further explored [67]. PHB2 and Nix are autophagy receptors located at the inner and outer mitochondrial membranes, respectively, and traditionally viewed as independent of the PINK1/Parkin pathway [68]. Our results suggest that Parkin expression correlates with PHB2 but not with Nix. During AAA development, PHB2 expression significantly increases while Parkin expression diminishes in AAA samples. In Parkin SMCKO AAA mice, there is a trend of increasing PHB2 expression in aortic tissues. Recent research shows that PHB2 is essential for Parkin-mediated mitophagy and conversely, Parkin can directly interact with and ubiquitinate PHB2 during mitophagy [69, 70]. Thus, our findings support a potential connection between PHB2 and the PINK1/Parkin mitophagy pathway, warranting further investigation into PHB2’s impact on PINK1/Parkin-mediated mitophagy during AAA.

Metabolomic analysis of Parkin SMCKO AAA serum reveals increased Acyl-C4-DC, a marker of deficient mitochondrial β-oxidation of fatty acids, potentially due to increased mitochondrial dysfunction in the AAA tissue of Parkin SMCKO mice. These data are consistent with previous findings that suggests synthetic VSMCs rely more heavily on fatty acid oxidation than glucose oxidation [71].Conversely, USP30 inhibitor treatment elevates serotonin levels in the serum. Recent studies indicate that serotonin enhances mitochondrial functions by increasing mitochondrial respiration and membrane potential, impacting reactive oxygen species and intracellular ATP [72]. Serotonin also regulates mitochondrial biogenesis via the SIRT1-PGC1α axis [73]. While the role of serotonin in AAA is not yet investigated, its competing pathway, the kynurenine pathway, is upregulated during AAA [74, 75]. Inhibition of Indoleamine 2–3 dioxygenase 1 (IDO), which converts tryptophan into the kynurenine pathway, has been shown to improve AAA outcomes [76]. This implies that upregulating the tryptophan-serotonin pathway might benefit AAA treatment, aligning with our findings that serotonin’s role in AAA merits further investigation.

USP30 antagonizes Parkin-mediated mitophagy, with studies showing that USP30 inhibition attenuates Parkinson’s disease, subarachnoid hemorrhage induced early brain injury, and kidney injury by inducing mitophagy [25, 77–79]; however, the role of USP30 in AAA remains largely unexplored. Our *in vivo* and *in vitro* experiments demonstrate that USP30 inhibitor treatment enhances Parkin-mediated mitophagy and PGC1α expression, promoting mitochondrial biogenesis, and significantly reducing AAA incidence. The potential mechanism behind promoting mitochondrial biogenesis could involve Parkin ubiquitinating PARIS, thus relieving its inhibition of PGC1α transcription [80]. This evidence supports the notion that USP30 inhibition and promotion of PINK1/Parkin-mediated mitophagy could be a promising therapeutic approach for AAAs.

Our study has several limitations. Treating aged AAA mice with USP30 inhibitors would enhance understanding, especially given the decreased baseline mitophagy levels in VSMCs from aged aortas. However, aneurysm incidence is very low when applying the same AngII model to aged mice, likely due to differences in aortic structure properties. Secondly, the designed intervention aims to prevent rather than treat potentially influencing clinical translation. Thirdly, the AngII and elastase AAA models do not fully represent human AAA pathogenesis and pathology; thus, using additional complementary AAA models could further validate our results and would represent potential future studies within this pathway. Future research could investigate potential regulatory mechanisms of Parkin and PINK1 in the context of AAAs. There is limited evidence on the upstream regulation of PINK1/Parkin, which remains to be explored. Research should also explore Parkin’s role in different cell types, as its function is suggested to extend beyond mitophagy pathways due to its ability to ubiquitinate additional targets.

In conclusion, our study reveals a decline in mitophagy within aortic VSMCs during aging and AAA formation, potentially linked to decreased PINK1/Parkin pathway activity. Parkin SMCKO induces exacerbated AAA formation, while utilizing USP30 inhibitor to enhance PINK1/Parkin-mediated mitophagy significantly reduces AAA incidence. The USP30 inhibitor, which also possesses therapeutic potential in other degenerative diseases, holds promise as a novel treatment for AAA.

## Supporting information

Manuscript text file

## Acknowledgements

The USP30i molecule VB-08 was provided at no cost by Vincere Biosciences Inc. (published in US Patent No. US11845724B2).

## Abbreviations

AAAs: abdominal aortic aneurysms
AngII: angiotensin II
mtDAMPs: mitochondrial damage – associated molecular patterns
EVAR: endovascular aneurysm repair surgery
VSMCs: vascular smooth muscle cells
mtDNA: mitochondrial DNA
HFD: high-fat diet
DMSO: dimethylsulfoxide
IP: intraperitoneal
MFI: mean fluorescence intensity
PKO: Parkin knock-out
TPM: transcripts per million
BP: biological process
CC: cellular component
MF: molecular function
DEG: differentially expressed genes
Parkin SMCKO: Parkin knock-out in smooth muscle cells
RBC: red blood cells
WT: wild-type
USP30i: USP30 inhibitor
shRNA: short hairpin RNA
SEM: standard error of the mean

