## Supplementary material for "Deficiency of mitophagy mediator Parkin in aortic smooth muscle cells exacerbates abdominal aortic aneurysm": Manuscript text file

**Supplementary materials**

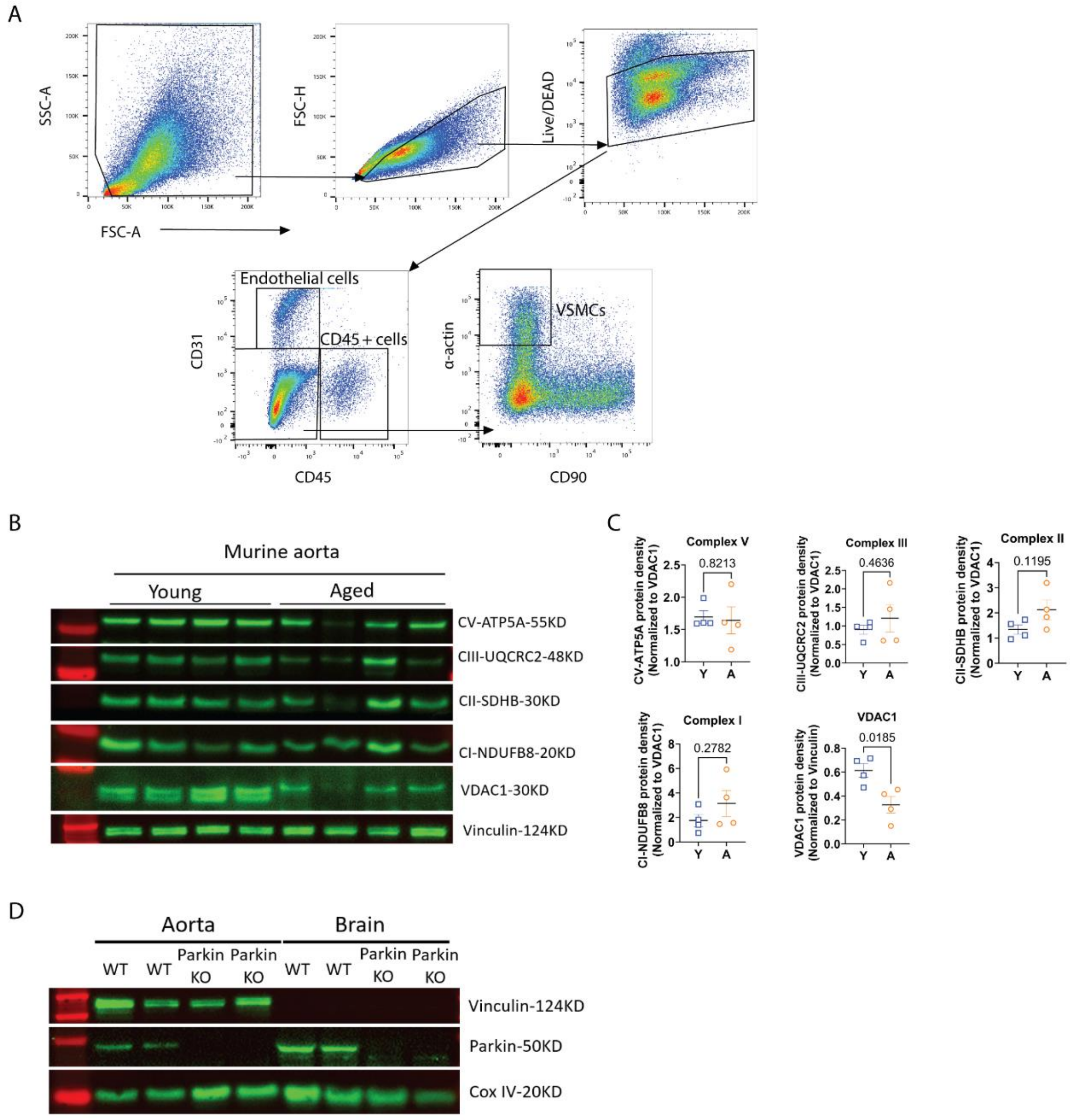


**Figure S1.** (A) Representative gating strategy for the identification of endothelial cells, CD45+ subsets and VSMCs from murine aorta. Following identification of cells by FSC-A vs SSC-A, doublets were excluded by FSC-A vs FSC-H, followed by exclusion of dead cells by live dead aqua. Endothelial cells were identified at $CD45^-$ and $CD31^+$. $CD45^+$ subsets were identified as $CD31^-$ and $CD45^+$ cellular populations. VSMCs were identified as $CD31^-$, $CD45^-$, $CD90^-$ and α-$actin^+$. (B-C) The contents of four representative proteins associated with mitochondrial complexes within the aortic tissues in Y and A group, including complex V - ATP5A, complex III - UQCRC2, complex II - SDHB, complex I - NDUFB8, were determined by Western blot, utilizing VDAC1 as internal control for quantification. (D) The validation of the specificity of Parkin antibody for Western blotting was conducted using protein extracts obtained from aortic and brain tissues isolated from both WT and Parkin global knockout mice. Results are presented as mean ± SEM. Unpaired two-tailed Student's t-test or Mann-Whitney U test were used for statistical analysis.

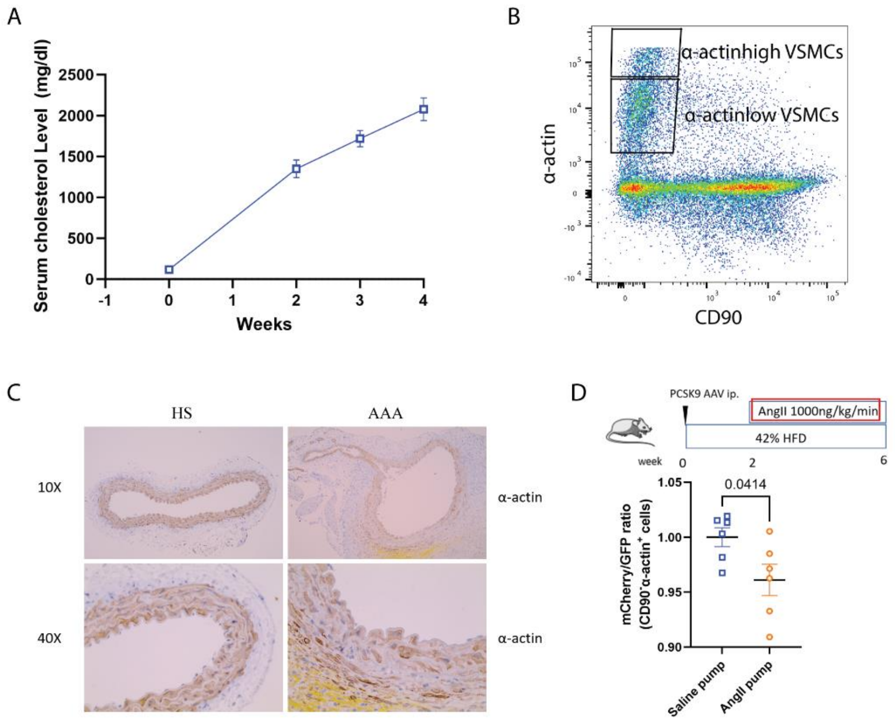


**Figure S2.** (A) Representative changes of serum cholesterol levels during the AAA development in the WT mice. (B) Representative adjusted gating strategy for VSMCs in the murine AAA. VSMCs were identified as $CD31^-$, $CD45^-$, $CD90^-$ and then classified as α-$actin^{high}$ VSMCs and α-$actin^{low}$ VSMCs by the expression amplitude of α-actin. (C) The representative aortic IHC staining images of α-actin in WT mice, HFD with Saline pump group versus HFD with AngII AAA group. Results are presented as mean ± SEM. Unpaired two-tailed Student's t-test or Mann-Whitney U test were used for statistical analysis. (D) Mice receiving 4-weeks AngII vs. Saline pump without HFD, aortas were harvested, digested, and stained with flow cytometric antibodies. Mitophagy levels of VSMCs (CD45-$CD90^-$$α^-$$actin^+$) were quantified by calculating the ratios of MFI of mCherry compared to GFP (n= 6 per group). Results are presented as mean ± SEM. Unpaired two-tailed Student's t-test or Mann-Whitney U test were used for statistical analysis.

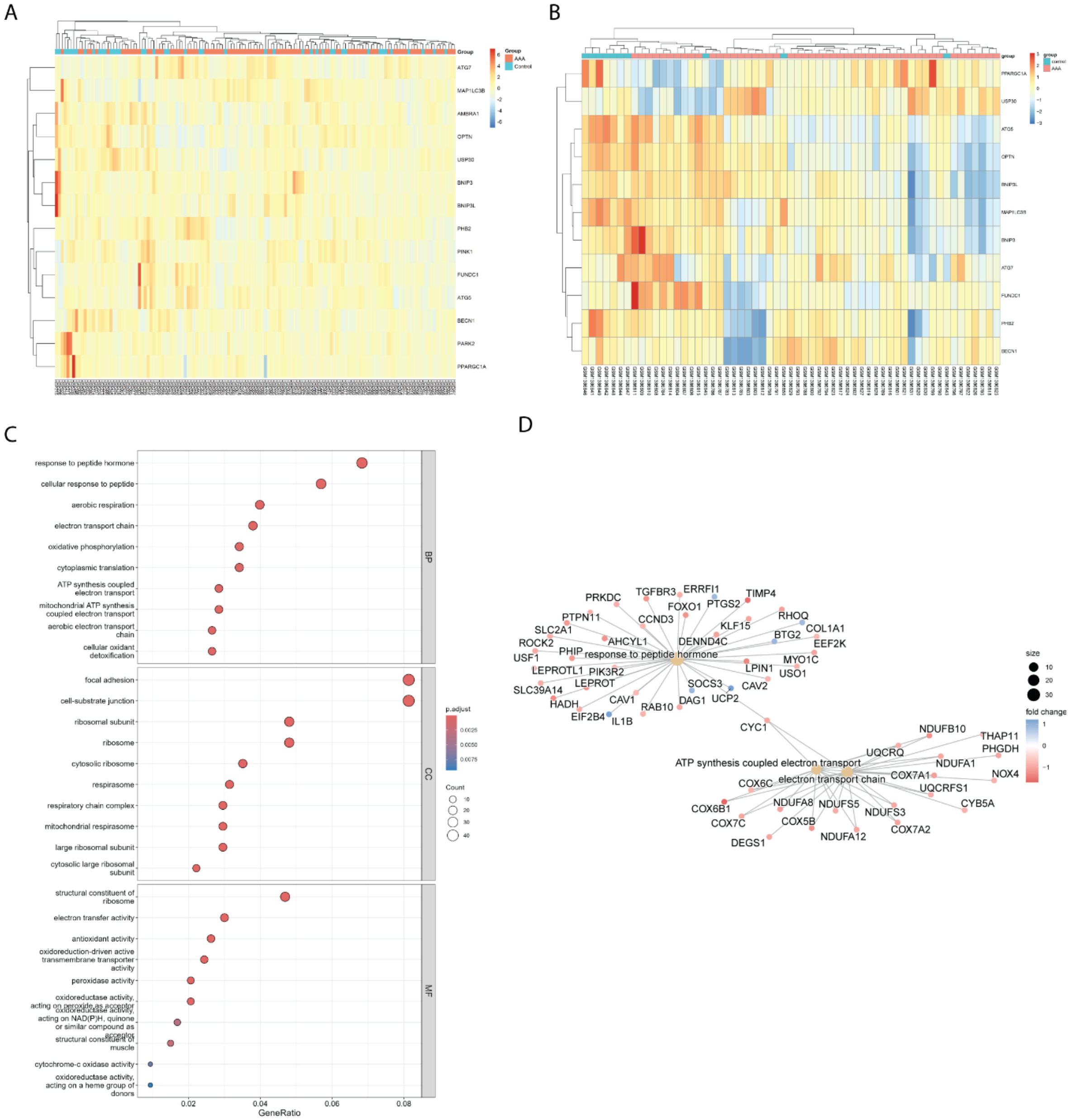
A
Group
AAA
Control
ATG7
MAP1LC3B
AMBRA1
OPTN
USP30
BNIP3
BNIP3L
PHB2
PINK1
FUNDC1
ATG5
BECN1
PARK2
PPARGC1A
B
group
control
AAA
PPARGC1A
USP30
ATG5
OPTN
BNIP3L
MAP1LC3B
BNIP3
ATG7
FUNDC1
PHB2
BECN1
C
response to peptide hormone
cellular response to peptide
aerobic respiration
electron transport chain
oxidative phosphorylation
cytoplasmic translation
ATP synthesis coupled electron transport
mitochondrial ATP synthesis coupled electron transport
aerobic electron transport chain
cellular oxidant detoxification
focal adhesion
cell-substrate junction
ribosomal subunit
ribosome
cytosolic ribosome
respirasome
respiratory chain complex
mitochondrial respirasome
large ribosomal subunit
cytosolic large ribosomal subunit
structural constituent of ribosome
electron transfer activity
antioxidant activity
oxidoreduction-driven active transmembrane transporter activity
peroxidase activity
oxidoreductase activity, acting on peroxide as acceptor
oxidoreductase activity, acting on NAD(P)H, quinone or similar compound as acceptor
structural constituent of muscle
cytochrome-c oxidase activity
oxidoreductase activity, acting on a heme group of donors
BP
CC
MF
p.adjust
0.0025
0.0050
0.0075
Count
10
20
30
40
0.02
0.04
0.06
0.08
GeneRatio
D
PRKDC
TGFBR3
ERRFI1
TIMP4
PTPN11
FOXO1
PTGS2
CCND3
RHOQ
SLC2A1
KLF15
COL1A1
ROCK2
AHCYL1
DENND4C
BTG2
EEF2K
USF1
PHIP
response to peptide hormone
MYO1C
LEPROTL1
PIK3R2
LPIN1
USO1
LEPROT
SOCS3
CAV2
SLC39A14
CAV1
DAG1
UCP2
HADH
RAB10
EIF2B4
IL1B
CYC1
NDUFB10
THAP11
UQCRQ
PHGDH
ATP synthesis coupled electron transport
NDUFA1
electron transport chain
COX7A1
NOX4
COX6C
UQCRFS1
COX6B1
NDUFA8
NDUFS5
NDUFS3
CYB5A
COX7C
COX5B
COX7A2
DEGS1
NDUFA12
size
10
20
30
fold change
1
0
−1

E

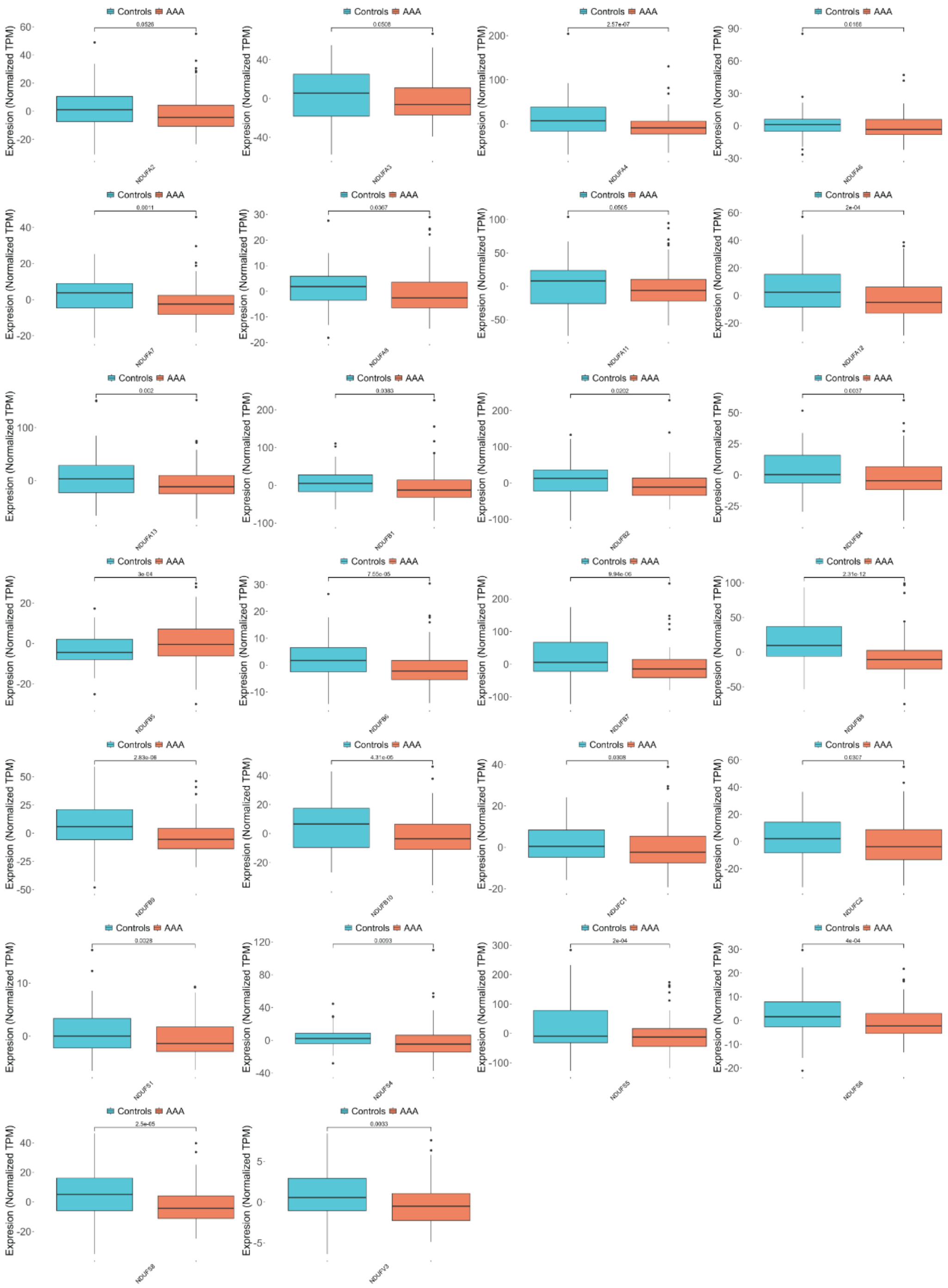
Controls AAA
Expresion (Normalized TPM)
0.0526
NDUFA2
0.0506
NDUFA3
2.57e-07
NDUFA4
0.0166
NDUFA6
0.0011
NDUFA7
0.0367
NDUFA8
0.0505
NDUFA11
2e-04
NDUFA12
0.002
NDUFA13
0.0383
NDUFB1
0.0202
NDUFB2
0.0037
NDUFB4
3e-04
NDUFB5
7.55e-05
NDUFB6
9.94e-06
NDUFB7
2.31e-12
NDUFB8
2.83e-06
NDUFB9
4.31e-05
NDUFB10
0.0308
NDUFC1
0.0307
NDUFC2
0.0028
NDUFS1
0.0093
NDUFS4
2e-04
NDUFS5
4e-04
NDUFS6
2.5e-05
NDUFS8
0.0033
NDUFV3

F

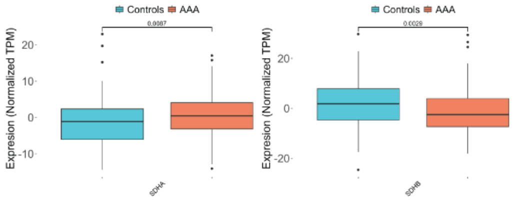
Controls AAA
Expresion (Normalized TPM)
0.0087
SDHA
Controls AAA
Expresion (Normalized TPM)
0.0029
SDHB

G

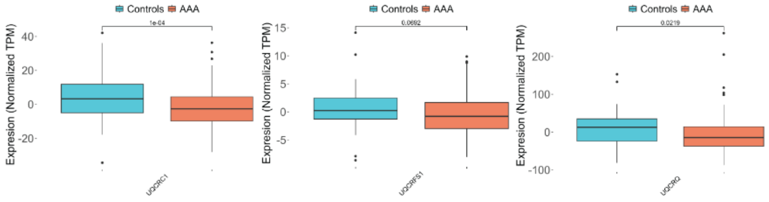
Controls AAA
Expresion (Normalized TPM)
1e-04
UQCRC1
Controls AAA
Expresion (Normalized TPM)
0.0692
UQCRFS1
Controls AAA
Expresion (Normalized TPM)
0.0219
UQCRQ

H

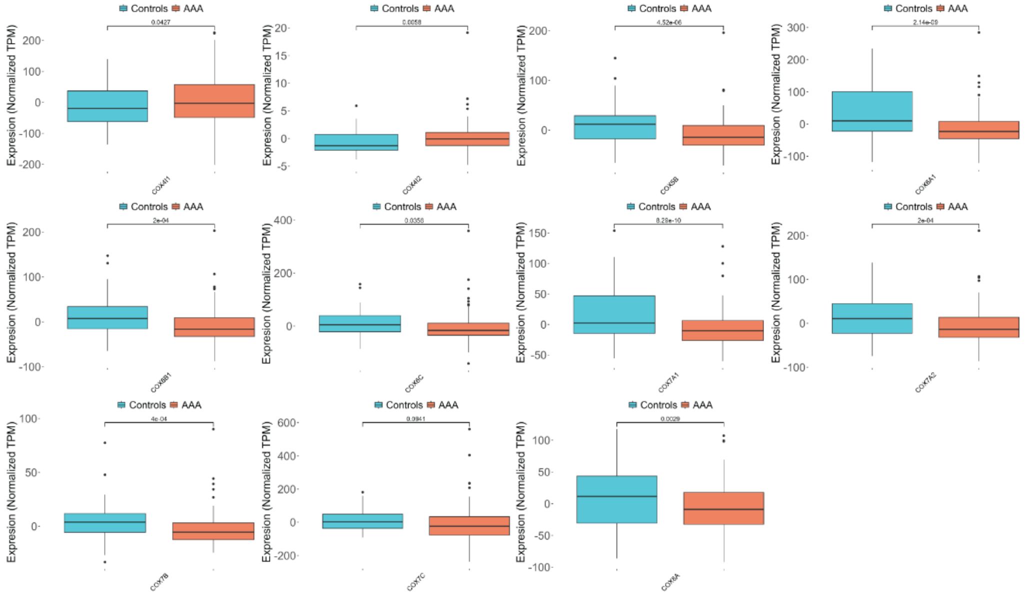
Controls AAA
Expresion (Normalized TPM)
0.0427
COX4I1
0.0058
COX4I2
4.52e-06
COX5B
2.14e-09
COX6A1
2e-04
COX6B1
0.0356
COX6C
8.28e-10
COX7A1
2e-04
COX7A2
4e-04
COX7B
0.0941
COX7C
0.0029
COX8A

I

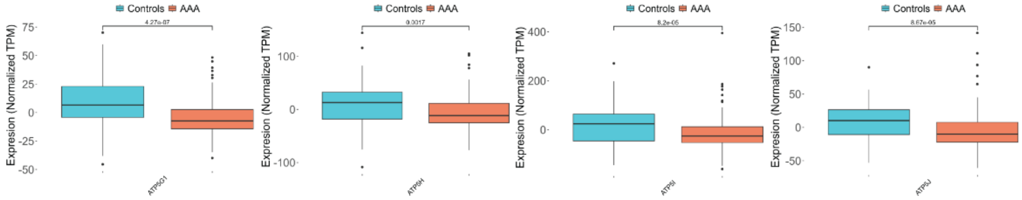
Controls AAA
Expresion (Normalized TPM)
4.27e-07
ATP5G1
0.0017
ATP5H
8.2e-05
ATP5I
8.67e-05
ATP5J

J

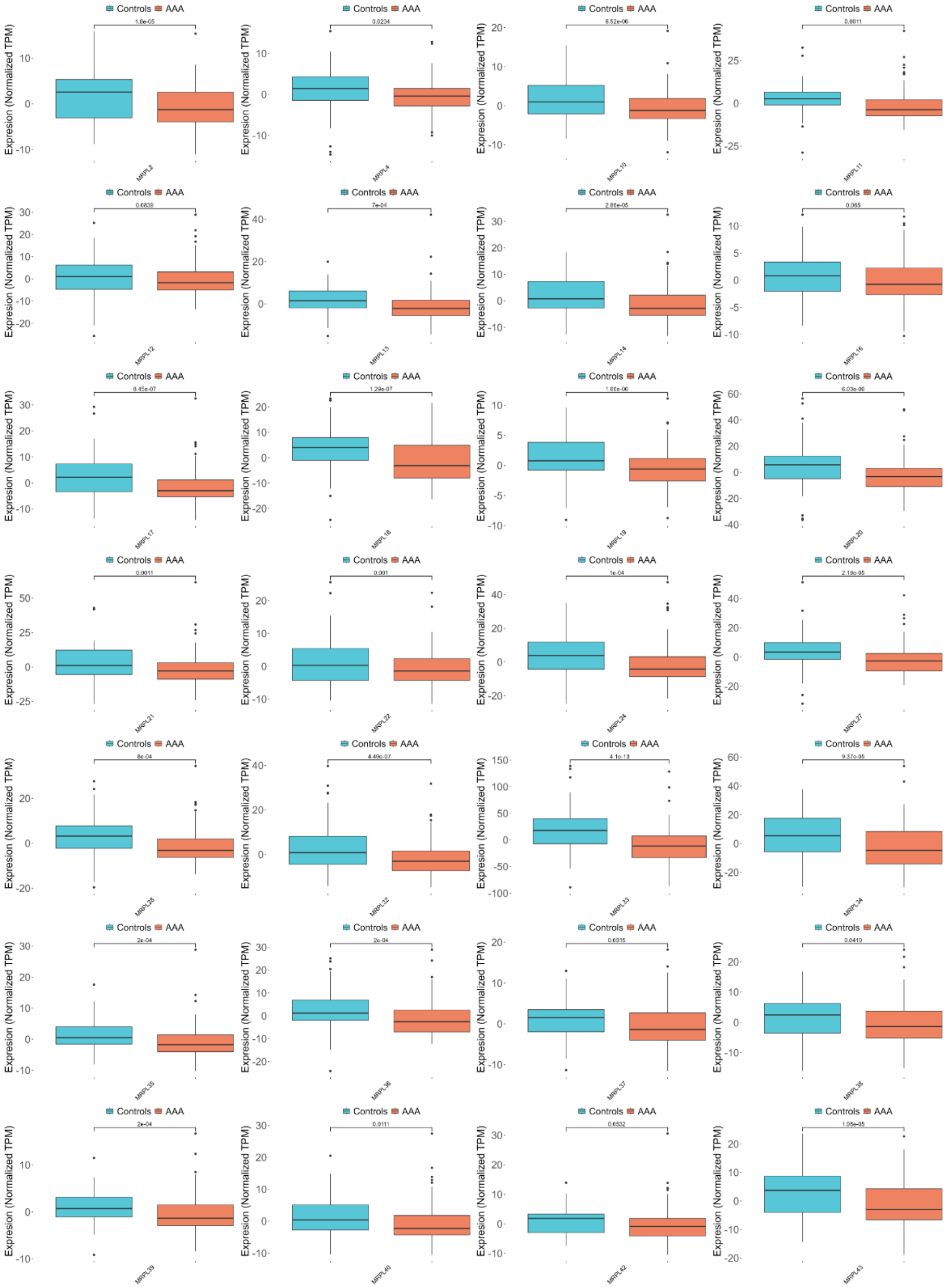

K

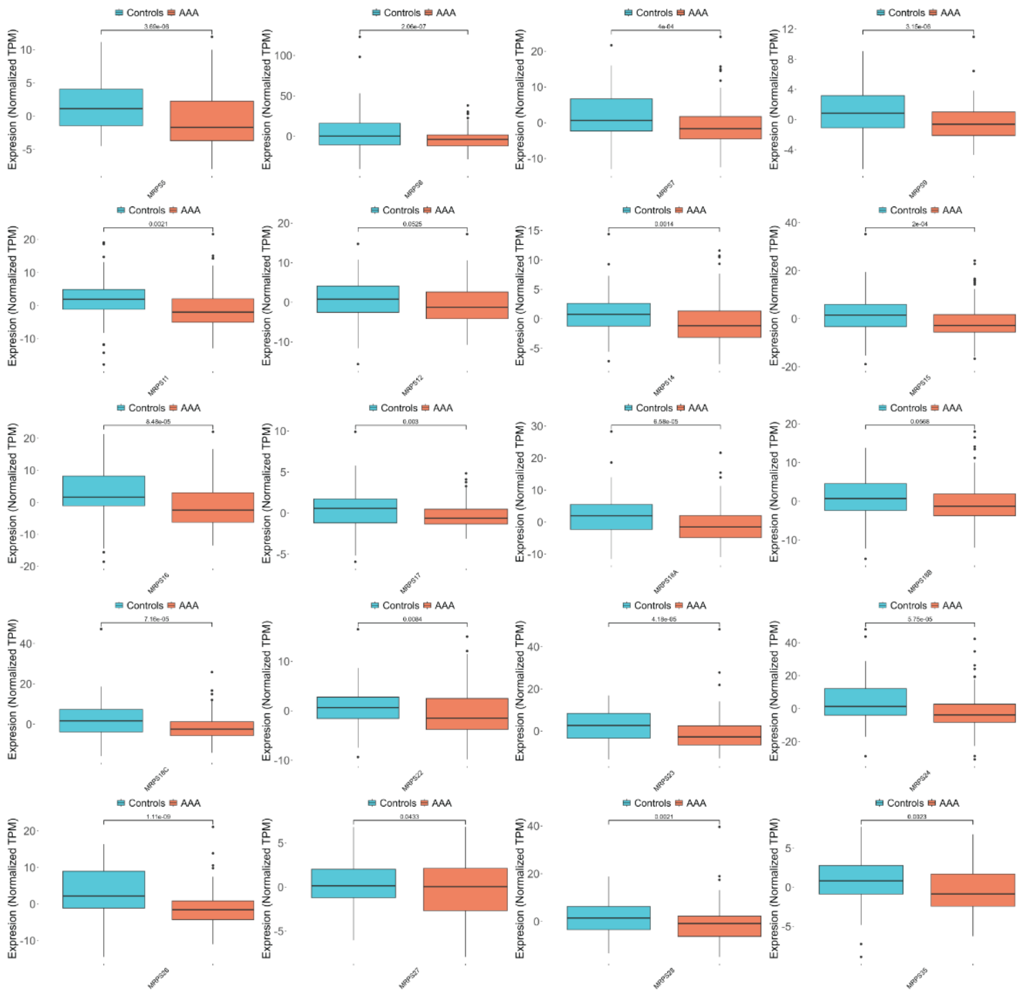


**Figure S3.** (A) Heatmap of gene expression clustered by mitophagy-associated genes in a human bulk RNA sequencing dataset (Control, n=44; AAA, n=96), highlighting PARK2, PINK1, BNIP3, BNIP3L, USP30, PHB2, ATG5, ATG7, MAPLC3B, AMBRA1, BECN1, FUNDC1, OPTN, and PPARGC1A. (B) Heatmap of gene expression clustered by PPARGC1A, USP30, ATG5, OPTN, BNIP3L, MAPLC3B, BNIP3, ATG7, FUNDC1, PHB2, and BECN1 from the publicly available bulk RNA sequencing dataset GSE57691. (C) KEGG pathway enrichment analysis of DEGs (differentially expressed genes) in dataset GSE57691, categorized by BP (biological process), CC (cellular component), and MF (molecular function). (D) Gene-concept network plot of the human bulk RNA sequencing dataset GSE57691 linking DEGs and BP terms. (E) Boxplots showing altered expression of nuclear genes encoding mitochondrial complex I in the human AAA bulk RNA sequencing dataset, including NDUFA2-8, NDUFA11-13, NDUFB1, NDUFB2, NDUFB4-10, NDUFC1, NDUFC2, NDUFS1, NDUFS4-6, NDUFS8, and NDUFV3. (F) Boxplots showing altered expression of nuclear genes encoding mitochondrial complex II in the human AAA bulk RNA sequencing dataset, including SDHA and SDHB. (G) Boxplots showing altered expression of nuclear genes encoding mitochondrial complex III in the human AAA bulk RNA sequencing dataset, including UQCRC1, UQCRFS1, and UQCRQ. (H) Boxplots showing altered expression of nuclear genes encoding mitochondrial complex IV in the human AAA bulk RNA sequencing dataset, including COX4I1, COX4I2, COX5B, COX6A1, COX6B1, COX6C, COX7A1, COX7A2, COX7B, COX7C, and COX8A. (I) Boxplots showing altered expression of nuclear genes encoding mitochondrial complex V in the human AAA bulk RNA sequencing dataset, including ATP5G1, ATP5H-J. (J) Boxplots showing altered expression of nuclear genes encoding the large subunit of the mitochondrial ribosome in the human AAA bulk RNA sequencing dataset, including MRPL2, MRPL4, MRPL10-14, MRPL16-22, MRPL24, MRPL27, MRPL28, MRPL32-40, MRPL42, and MRPL43. (K) Boxplots showing altered expression of nuclear genes encoding the small subunit of the mitochondrial ribosome in the human AAA bulk RNA sequencing dataset, including MRPS5-7, MRPS9, MRPS11, MRPS12, MRPS14-17, MRPS18A, MRPS18B, MRPS18C, MRPS22-24, MRPS26-28, and MRPS35.

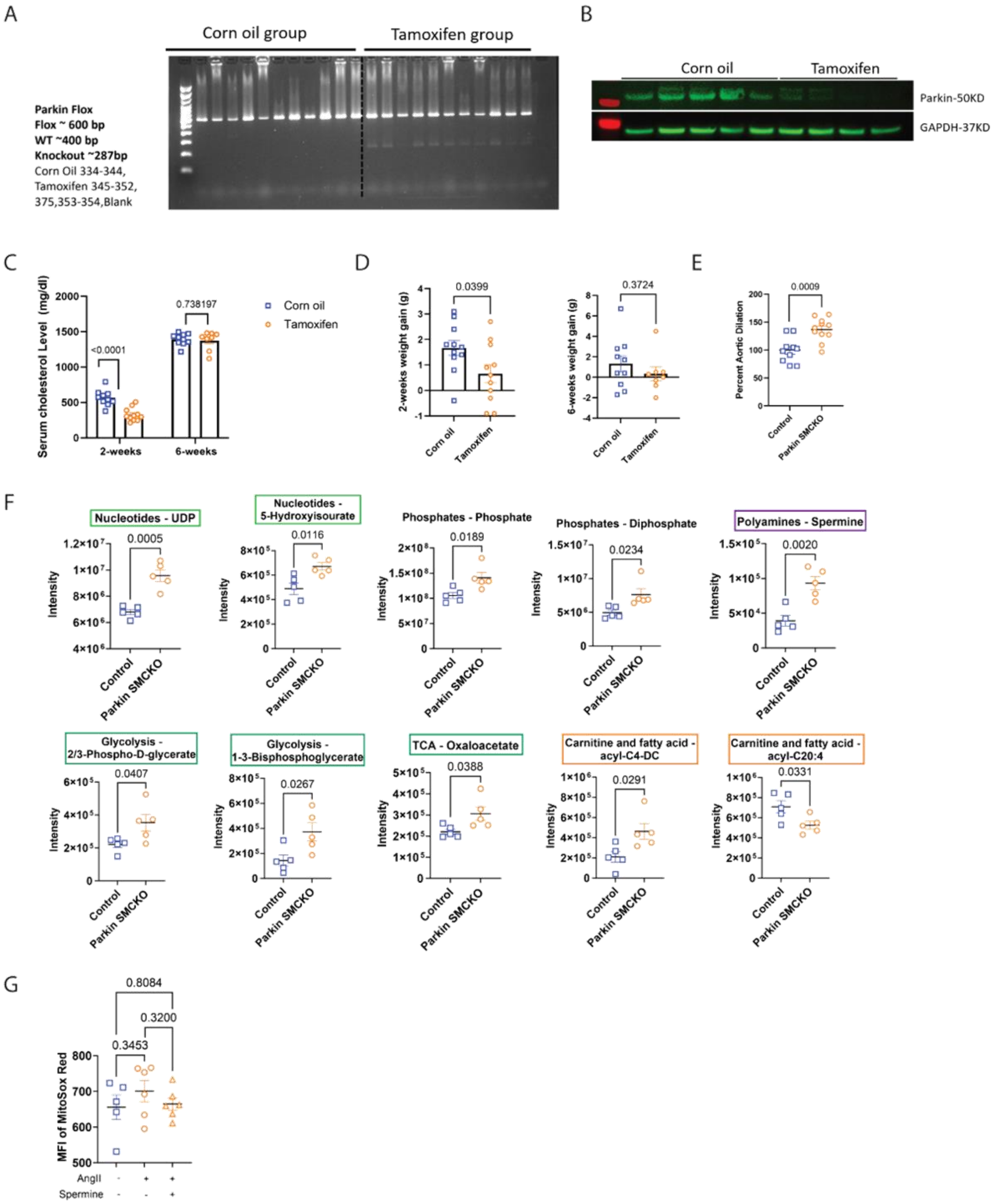


**Figure S4.** (A) Genotyping results of Myh11-creERT2 Parkin fl/fl mice, which were administered corn oil or tamoxifen (n=11 per group). (B) Parkin expression level in the descending aortas of corn oil and tamoxifen treated Myh11-creERT2 Parkin fl/fl mice, were determined by Western blot (n=4-5 per group), employing GAPDH as internal control for quantification. (C-D) Changes in body weight throughout the AAA development within the Myh11-creERT2 Parkin fl/fl mouse cohort were measured and recorded. (E) Aortic dilation between control and Parkin SMCKO group in the elastase AAA model. (F) Metabolites from different metabolomic pathways that have significant alterations between control and Parkin SMCKO groups. (G) The MFI of MitoSox Red (n=5-6 per group) within MOVAS respectively treated by vehicle, 100nM AngII and a combination of 100nM AngII with 100nM spermine, were quantified and showed in histogram. Results are presented as mean ± SEM. Unpaired two-tailed Student's t-test or Mann-Whitney U test were used for statistical analysis.

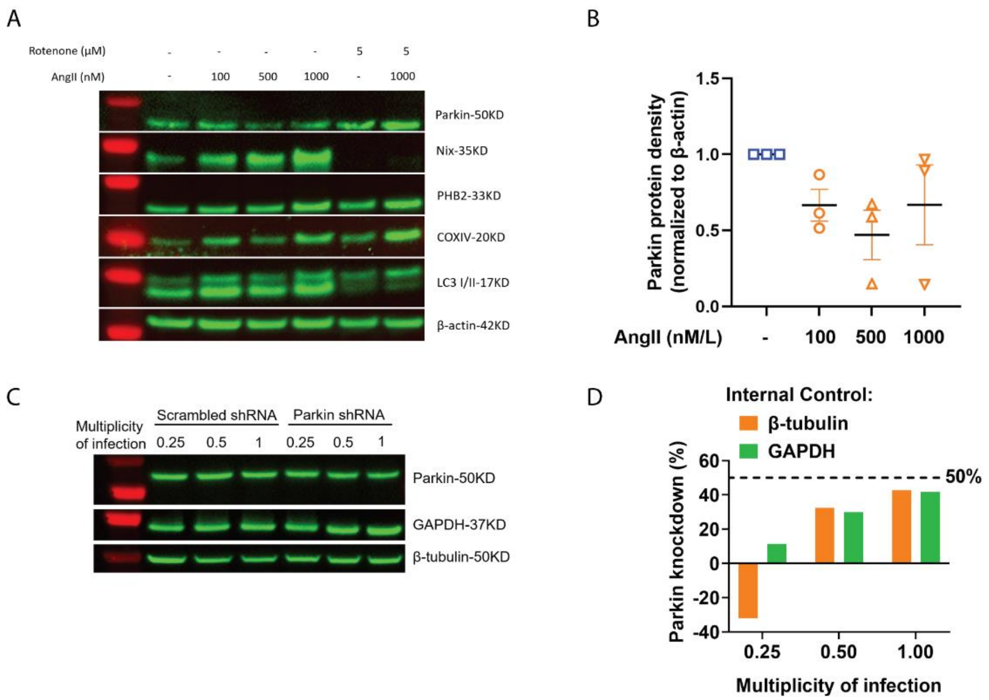


**Figure S5.** (A-B) The AngII dose titration to treat MOVAS was performed to assess its impact on the expression of relevant proteins, including Parkin, Nix, PHB2, COXIV, LC3I/II. The quantification of Parkin expression is displayed (n=3 per group). (C-D) Knock down of Parkin at multiplicity of infection as 0.25, 0.5 and 1 by lentiviral Parkin shRNA was quantified by Westernblot.

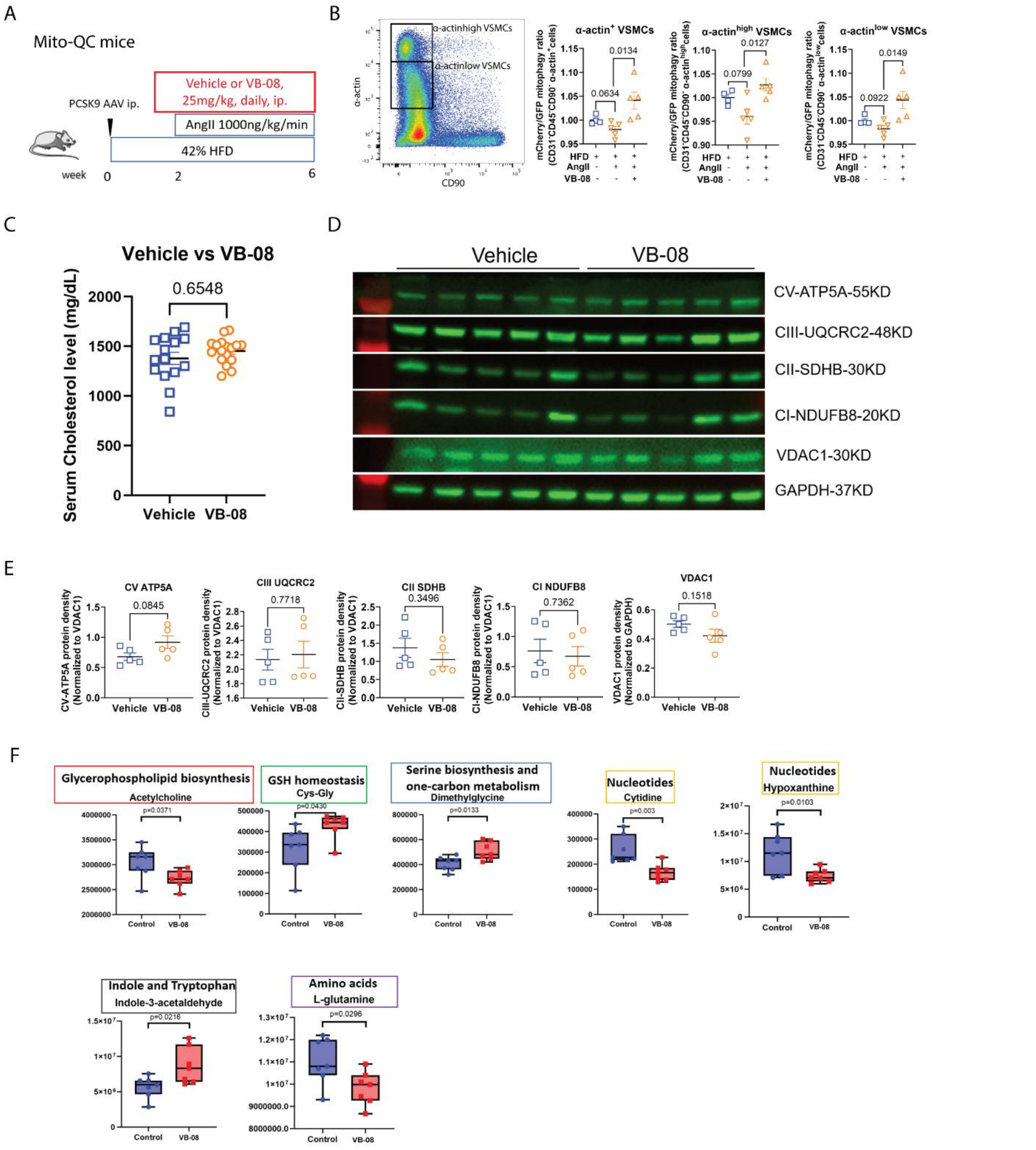


**Figure S6.** (A) The Scheme of employing Mito-QC mice to investigate VB-08's impact on the AAA model. (B) Pseudo color plot showing the gate of VSMCs – CD90$^{-}$α-actin$^{high}$ and CD90$^{-}$α-actin$^{low}$ populations, mitophagy levels of which were quantified by calculating the ratios of mean fluorescence intensity (MFI) of mCherry versus GFP across three groups: HFD + Saline pump + Vehicle, HFD + AngII pump + Vehicle and HFD + AngII pump + VB-08. (C) Measurement of serum cholesterol levels in C57BL/6 mice cohort after 6 weeks, comparing the vehicle treated to the VB-08 treated group. (D-E) The contents of four representative proteins associated with mitochondrial complexes within the descending aortic tissues in vehicle and VB-08 group, including complex V - ATP5A, complex III - UQCRC2, complex II - SDHB, complex I - NDUFB8, were determined by Western blot, utilizing VDAC1 as internal control. (F) Metabolites from different metabolomic pathways that have significant alterations between vehicle and VB-08 treated groups. Results are presented as mean ± SEM. Unpaired two-tailed Student's t-test or Mann-Whitney U test were used for statistical analysis.
